# Biodiversity is enhanced by sequential resource utilization and environmental fluctuations via emergent temporal niches

**DOI:** 10.1101/2023.02.17.529002

**Authors:** Blox Bloxham, Hyunseok Lee, Jeff Gore

## Abstract

How natural communities maintain their remarkable biodiversity and which species survive in complex communities are central questions in ecology. Resource competition models successfully explain many phenomena but typically predict only as many species as resources can coexist. Here, we demonstrate that sequential resource utilization, or diauxie, with periodic growth cycles can support many more species than resources. We explore how communities modify their own environments by sequentially depleting resources to form sequences of temporal niches, or intermediately depleted environments. Biodiversity is enhanced when community-driven or environmental fluctuations modulate the resource depletion order and produce different temporal niches on each growth cycle. Community-driven fluctuations under constant environmental conditions are rare, but exploring them illuminates the temporal niche structure that emerges from sequential resource utilization. With environmental fluctuations, we find most communities have more stably coexisting species than resources with survivors accurately predicted by the same temporal niche structure and each following a distinct optimal strategy. Our results thus present a new niche-based approach to understanding highly diverse fluctuating communities.

## Introduction

From coral reef fishes and grassland plants to ocean plankton and gut microbiota, natural communities display remarkable biodiversity. Yet, how this diversity is stably maintained has long puzzled researchers [1–9]. As all organisms must consume resources to grow and survive, resource competition is often assumed to be a dominant interspecies interaction [2,10–13]. However, the simplest formulations of resource competition predict a “competitive exclusion principle” that there should not be more surviving species than resources to compete for [3,4,10,14]. Non-equilibrium temporal dynamics, in the form of internally driven oscillations [15,16], external environmental fluctuations [17,18], or seasonal cycles [19], were an early suggestion for stabilizing more coexisting species than resources [4,20] and have now been established as a key explanation for the biodiversity observed in nature [5,6,21–33]. The importance of temporal dynamics in sustaining biodiversity highlights the need to thoroughly understand the competitive interactions and ecological niches that emerge in fluctuating communities.

Multiple studies have demonstrated that fluctuations in species abundance can increase community diversity [21,27,30,34–36], although the role of resource competition in fluctuating communities has only been explored in a few specific contexts [37]. Huisman et al. demonstrated in a model of simultaneous consumption of multiple essential resources that random communities will often reach a chaotic attractor or limit cycle in which more species than resources coexist [6,22,23,38,39]. Interestingly, this diversity was not enabled by producing an enlarged set of distinct ecological niches but instead by placing species on a continuum in which another species with intermediate characteristics could always invade the community [23,38]. Thus, species do not play clearly distinct roles in the community, coexistence is often neutral or transient rather than truly stable [23,37], and it remains unclear how resilient these communities and their diversity are to perturbations [23,40,41]. Recently, Sakavara et al. demonstrated that adding a fluctuating resource supply to this model led to the robust clustering of species (a.k.a. “lumpiness” [42]) based on relative investment in different resources. Intriguingly, these clusters often outnumber the resources, offering fresh perspectives on how diverse communities organize themselves according to resource competition strategies [37]. Meanwhile, Pacciani-Mori et al. showed that if species can adapt their metabolic strategies while subject to a strict metabolic tradeoff then highly diverse communities are possible under either a constant or periodically oscillating resource supply [43]. However, it remains unclear whether highly diverse communities are likely to coexist when other forms of resource competition dominate. Exploring other forms of resource competition may also further illuminate the distinct roles that different species play in community dynamics.

Previous consideration of resource competition has typically assumed that species simultaneously utilize all resources available to them [10,22,37,44,45]. However, in nature sequential utilization of resources, also known as diauxie in the case of microbial growth, is a ubiquitous phenomenon [43,46–59]. For example, a microbe might consume glucose first, lactose only after glucose depletion, and acetate last when all three are initially available. Even in complex environments microbes initially consume only a subset of the available resources and consume the remaining resources only after the first set has been depleted [53]. Sequential resource utilization is also observed in macroscopic ecosystems, such as when foraging animals search for lower preference resources only when others are unavailable [47,51,54]. Some recent works have used diauxie as a lens for understanding and predicting microbial community assembly [59], including how the resource preferences of coexisting species relate to each other [57], while other work has explored how the structure of central metabolism affects the relative frequency of different preference orders emerging [60]. However, scientific understanding of how sequential utilization affects community diversity and dynamics remains limited.

Because sequential resource utilization implies the sequential depletion of resources, it naturally creates distinct temporal niches between each resource depletion. Temporal niches are a frequently invoked concept when considering seasonal and other fluctuating environments [29,61–67]. Whereas “niche” most commonly refers to a resource a species consumes, “temporal niche” refers to a period of time in which a species has some specific fitness, for example a time of day or a season. In this paper, “temporal niche” will refer to a period of time in which a given set of resources are available. Just as a species may compete for many resources, a species may be active in multiple temporal niches but with a different fitness in each. Under stochastic fluctuations temporal niches do not need to occur periodically but can instead be the periods of time when the system happens to be in a particular state given the fluctuations – a concept that will be explored in this paper.

Here, we use a simple model of sequential resource utilization under periodic cycles of growth, death, and resource resupply to demonstrate the coexistence of many more species than resources due to temporal fluctuations and illuminate the temporal niche structure of these communities. We first consider community-driven oscillations under constant environmental conditions to develop our understanding of the temporal niche structure, then consider stochastically fluctuating environments, and finally seasonal cycles. Community-driven oscillations (in which inter-species competition drives oscillations in community composition with periods longer than the growth cycles) are rare, but when they do occur exponentially many species as resources can coexist. This enhanced coexistence is possible because fluctuating population sizes drive variations in the resource depletion order and therefore variations in which temporal niches (the periods of time in which sets of resources are available) occur on each growth cycle. We then introduce extrinsic environmental fluctuations in the form of a stochastically fluctuating resource supply and observe that, with even small fluctuations, the vast majority of communities violate the competitive exclusion principle. We show that the same temporal niche structure applies to both community-driven oscillations and environmental fluctuations and use our understanding of temporal niches to accurately predict which species survive. We also show similar results under seasonal fluctuations. Our work thus builds upon previous understanding of highly diverse communities by demonstrating their emergence under very different forms of resource competition and by elucidating the temporal niches that define their competitive structure.

## Results

### Sequential utilization supports highly diverse communities when oscillations occur

To better understand how sequential resource utilization dictates community structure and dynamics, we developed a simple model based on exponential growth and a pulsed resource supply, capturing, for example, seasonal resource availability or a serial dilution experiment. We assumed the period of the supply pulses was long enough that species completely depleted one pulse of resources before the next, creating a series of discrete growth cycles (although future research should explore what happens when resources are not fully depleted by the next supply pulse, which could be a better model for some ecosystems). In the case of a single resource, species grow at constant exponential rates until the resource is depleted, at which point the population sizes are divided by 10 (representing death during the starvation period), the resource concentration is returned to its supply value, and another growth cycle begins (Fig 1A, Methods). Because species have a single, constant growth rate with only one resource, the same species will always be the fast-grower and will outcompete all others (Fig 1B). Thus, only a single species survives on a single resource.

**Figure 1.**
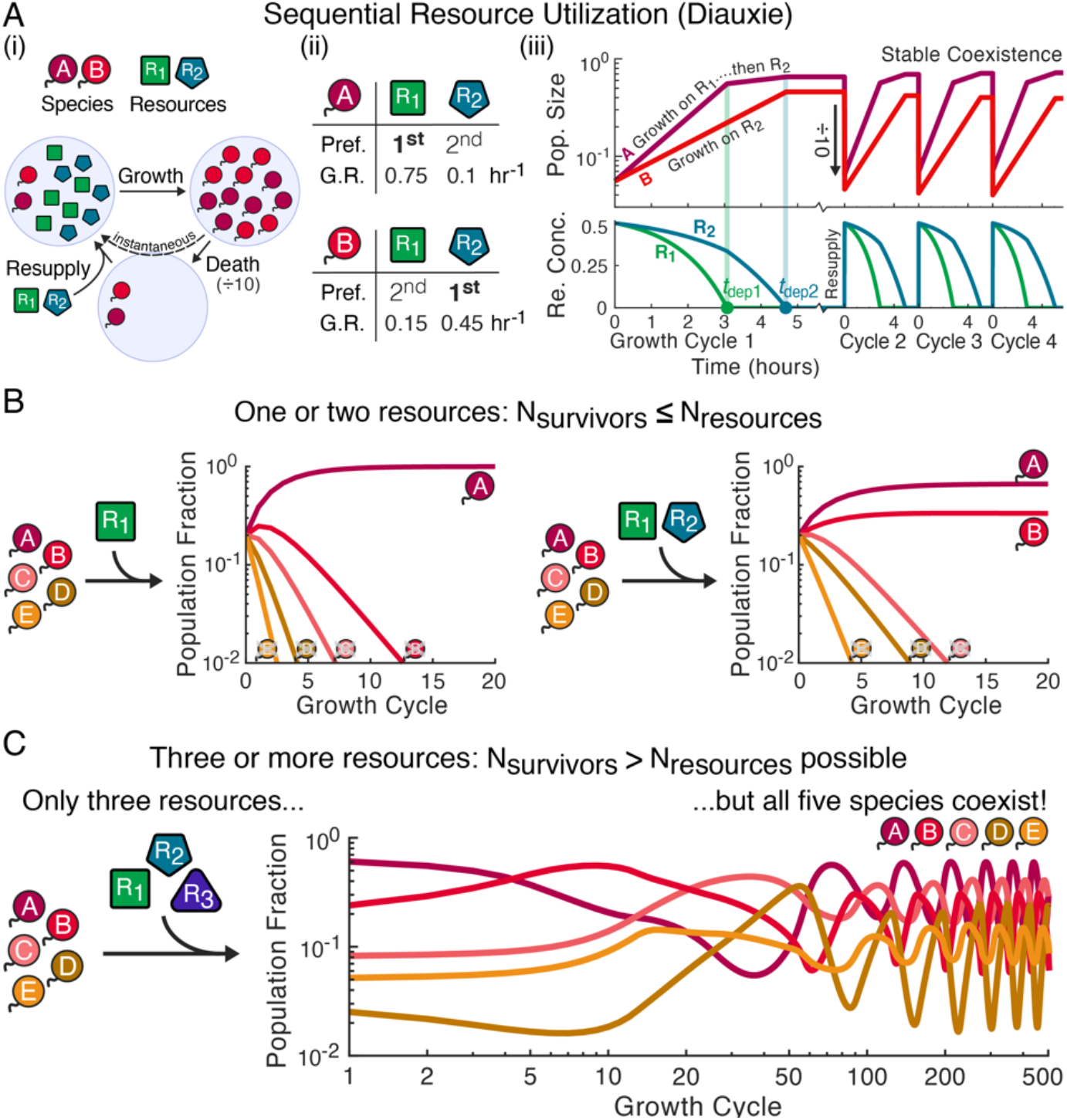
Diauxic resource competition can violate the competitive exclusion principle due to community-driven oscillations. (**A**) Definition of our model of resource competition with sequential utilization, also known as diauxie. (i) We assume a model of boom– and–bust cycles. Throughout all figures, circles with tails represent species and polygons represent resources. Species are inoculated into a well-mixed environment with finite quantities of resources available and fully deplete those resources as their populations grow. Species then experience a mortality before an instantaneous influx of resources and the start of the next growth cycle (Methods). (ii) Species are defined by their resource preferences orders (Pref.) and growth rates (G.R.) for each resources (Methods), as is illustrated for two example species. (iii) Simulated growth (top) and resource consumption (bottom) dynamics for those two species over four growth cycles. The resources depletion times determine each species’ overall growth on each cycle and are indicated with vertical lines. Species A and B coexist because, while A initially outpaces B, after R_1_ is depleted A’s growth slows while B continues growing at its maximum growth rate. (**B**) When species compete for only one or two resources, the competitive exclusion principle, which predicts that only as many species as resources can survive, is obeyed. Species’ population fractions at the end of each growth cycle for two example communities are shown (Appendix §1.1 for parameters). (**C**) However, when three resources are supplied, diauxie can produce community-driven oscillations and competitive exclusion violations. Shown is an example of five species competing for three resources with all five stably coexisting (Appendix §1.1 for parameters and Figure 2 for further exploration of this example).

With two or more resources, new dynamics arise because species now have ordered resource preferences that can vary between species and growth rates for each resource (Methods). Figure 1A illustrates an example in which species A prefers resource R_1_ over R_2_ while species B prefers R_2_ over R_1_. Species A is the initial fast-grower because its top-preference growth rate is faster than B’s. However, A’s faster growth means R_1_ is depleted before R_2_ and A needs to switch to R_2_. In the example, A’s R_2_ growth rate is much slower than B’s, and B is able to catch back up to A before R_2_ is also depleted (Fig 1A). Feedback between the resource depletion times and species’ population sizes allows the two species to stably coexist, converging towards a fixed point with constant population sizes from one growth cycle to the next (Fig 1A-B, Appendix §2.1). In the Appendix we show that when competing for two resources a fixed point is always reached (Appendix §3.1) and, with any number of resources, if a fixed point is reached there can be no more stably coexisting species than resources (Appendix §2.2). Thus, with one or two resources, at most one species per resource can stably coexist, obeying the “competitive exclusion principle” (Fig 1B).

However, with three or more resources, we discovered communities that did not reach fixed points but instead oscillated with periods longer than that of the growth cycles and had more surviving species than resources (Fig 1C). These examples were rare when sampling random communities (Appendix §3.2), but we were able to construct plentiful examples by choosing species likely to oscillate, confirming those oscillations, and adding additional species that could coexist with the original (Appendix §3.6 for more detail on this process and an intuitive construction of one such community). The example in Figures 1C and 2 has “anomalous” resource preferences [57] that do not correlate to a species’ growth rate order, but these are not necessary for oscillations to occur (Appendix §3.3). This example oscillation was robust to demographic noise (Appendix §3.7). In addition to periodic oscillations, chaos was also observed (Appendix §3.4), with chaotic communities also able to violate competitive exclusion. It appears larger communities were more likely than smaller ones to be chaotic, but our constructive approach did not allow us to meaningfully quantify or verify this trend. Such rich dynamics and large increases in diversity suggested a valuable opportunity for studying and understanding highly diverse, resource competition-driven communities.

### Temporal niches explain how fluctuations increase diversity and predict exponentially many species as resources could coexist

Having demonstrated that communities of sequential utilizers can maintain high diversity via spontaneous fluctuations, we proceeded to study the niche structure of these communities. Fluctuating population sizes cause resource depletion times to also fluctuate, and, as these depletion times are fundamental to interspecies interactions (Appendix §2.1), we began by looking at how they varied during oscillations. Focusing on the community of five species on three resources originally presented in Figure 1C (Fig 2A), we explored the growth and resource consumption dynamics on different growth cycles (Fig 2B).

**Figure 2.**
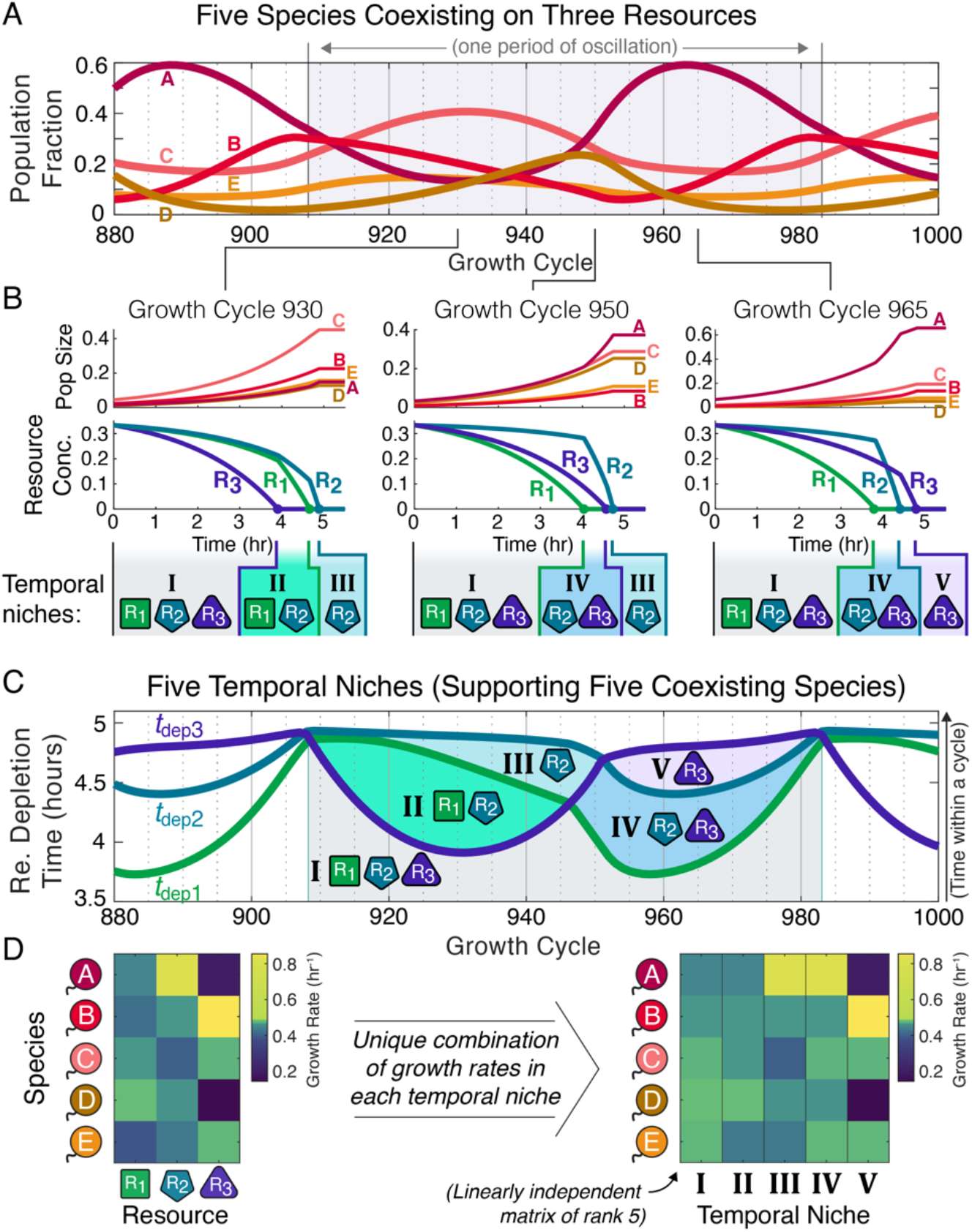
Oscillations allow for highly diverse communities due to an emergent temporal-niche structure with more distinct niches than resources. (**A**) Population fractions at the end of each growth cycle for the example of five species coexisting on three resources presented in Figure 1C (Appendix §1.1 for parameters). (**B**) Growth and resource-consumption dynamics on three different growth cycles. Top row shows population growth over the course of each growth cycle, while middle row shows decaying resource concentrations. Because each resource is depleted at a different time, species grow first in the three-resource environment, then in a two-resource environment, and finally in a single-resource environment. These sequentially realized environments are “temporal niches”. Fluctuating population sizes cause resources to be depleted in different orders on different growth cycles, so which temporal niches occur also varies, as is highlighted in the bottom row. (**C**) Resource depletion times (with *t*_dep *i*_ being the time spent in a cycle until resource R_*i*_ is depleted) across the entire period of the oscillation show a total of five temporal niches. Lines show the resource depletion times and shaded regions highlight the temporal niches. (**D**) Species’ growth rates by resource (left) and in each temporal niche (right). Species only have one growth rate per resource, but their differing resource preferences produce different combinations of growth rates in each temporal niche such that each niche becomes a distinct growth phase with independent dynamics.

We immediately noted that not only did the depletion times fluctuate, but the fluctuations were large enough that the order in which resources were depleted varied between cycles (Fig 2B). This varying depletion order has important consequences. On each growth cycle species grow in an increasingly depleted environment, first with all three resources present, then with all but one, and eventually on a single resource (Fig 1B and 2B). We refer to these sequential environments as “temporal niches”. In an oscillation with a variable resource depletion order, different sets of temporal niches occur on each growth cycle (Fig 2B). In the Figure 2 example, on Cycle 930 of the simulation species grow first in the all-resource niche then in the R_1_-and-R_2_ niche and finally in the R_2_-only niche (Fig 2B). However, by Cycle 950, R_1_ is depleted before R_3_ (Fig 2B) due to the rising population fraction of species A, which prefers R_1_, and declining fraction C, which prefers R_3_ (Appendix §1.1). Because R_1_ is now depleted before R_3_, the two-resource temporal niche is now the R_2_-and-R_3_ niche (Fig 2B). Considering the resource depletion times across the entire oscillation, we tallied five temporal niches (Fig 2B and 2C) – five niches allowing five species to coexist on only three resources.

Although species have fixed growth rates for each resource and consume only one resource at a time, each temporal niche has distinct dynamics: species’ growth rates are distributed across the temporal niches according to their resource preference orders such that no temporal niches share the same combination of species’ growth rates (Fig 2D and Appendix §2.3). Thus, the environmental mediation of community interactions has enough degrees of freedom that conditions exist in which as many species as temporal niches can exactly match the periodic mortality and therefore coexist (Appendix §2.3). Differing resource preferences also create rich patterns in when and how often species are direct competitors and to which depletion times each species contributes on each growth cycle, allowing for the necessary independent degrees of interaction to stabilize coexistence of the community (Appendix

§2.3). Temporal niches thus explain how highly diverse communities can arise from resource competition when temporal fluctuations occur. Notably, the temporal niches that arise under diauxic resource competition are a discrete set – as opposed to the continuum observed in previous models of highly diverse, fluctuating communities [23]. This property allows for calculating bounds on the number of coexisting survivors, sets of distinct optimum strategies, and which species are the most direct competitors, as is explored in the rest of this paper.

There can be one coexisting species for each temporal niche, and the maximum number of temporal niches is the number of combinations of whether each resource has been depleted yet, excluding the case of all resources being depleted (Fig 3A). This sets 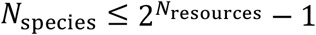 as an upper bound on the number of coexisting species, and suggests that exponentially many species as resources may be able to coexist when oscillations or chaos occur (Fig 3B; Appendix §2.3). We are not aware of a proof that this bound is achievable for arbitrary numbers of resources, but we leveraged our knowledge of the temporal niche structure (Appendix §2.3) to construct and simulate communities in which 7 species coexisted on 3 resources and 15 coexisted on 4 resources, which each saturated the upper bound, as well as examples of 23 species on 5 resources, which saturated 74% of the upper bound (Appendix §3.5), although increasing equilibration times to definitively demonstrate community stability made finding larger communities impractical. It therefore appears likely that exponentially many species as resources can coexist under diauxic resource competition when community-driven oscillations or chaos occur.

**Figure 3.**
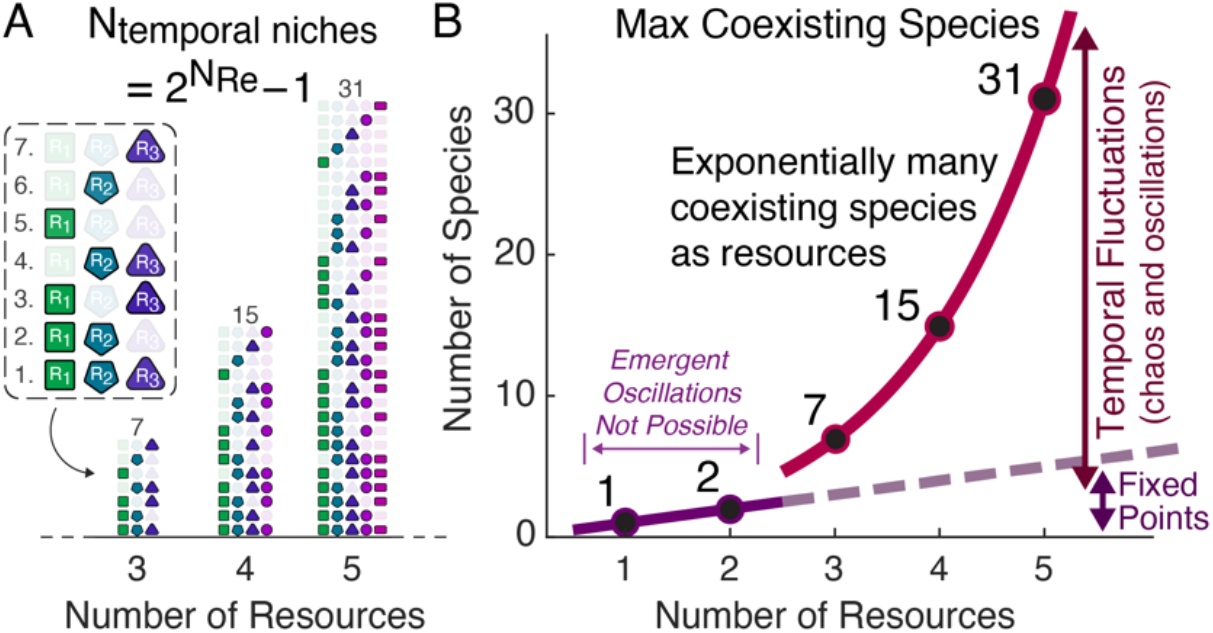
The maximum number of coexisting species scales exponentially with the number of resources. (**A**) The number of possible temporal niches is the number of binary combinations of whether or not each resource is still present 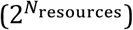 minus the combination corresponding to all resources being depleted. Shown are all the possible temporal niches for the cases of three, four, and five resources. (**B**) There can be at most one surviving species for each temporal niche, creating an upper bound on the maximum number of coexisting species that rises exponentially with the number of resources, as shown here in red. For comparison, one species per resource (the traditional competitive exclusion principle) is shown as a dashed gray line. Oscillations or chaos are necessary for more species than resources to coexist and require at least three species and three resources (Appendix §3.1), so the bound of one species per resource applies if there are only one or two resources (blue line).

### Most random assemblages have more coexisting survivors than resources in the presence of environmental fluctuations

Having established that having more coexisting species than resources (i.e. competitive exclusion violations) was possible with a constant resource supply, we proceeded to consider a more realistic case in which the resource supply is not strictly constant but instead fluctuates over time. When constructing examples with a constant resource supply, we had noted that community-driven oscillations, and therefore competitive exclusion violations, were rare (Appendix §3.1). As perturbations are often destabilizing, one might expect competitive exclusion violations under diauxic growth to become even less likely in the presence of environmental fluctuations. However, the competitive-exclusion violations we have demonstrated depended primarily on the resource depletion order varying, and environmental fluctuations would drive depletion-time fluctuations, so we speculated that the frequency of competitive exclusion violations and community diversity might actually increase in the presence of environmental fluctuations.

To test whether environmental fluctuations would indeed increase the frequency of competitive exclusion violations, we defined sampling distributions for resource supply on each growth cycle as a function of the environmental fluctuation magnitude σ_RS_ (Fig 4A-B, Methods). The fluctuation magnitude varied from a constant resource supply at σ_RS_ = 0 to a uniform-random supply at σ_RS_ = 0.236 (Fig 4B). We generated 100 communities each containing 5000 random species with the metabolic constraint that the L2-norm of each species’ growth rates equaled 1 hr^-1^ (i.e. 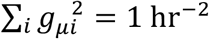 where *g*_*μi*_ is the growth rate of species μ on resource R_*i*_), simulated these communities at each of 9 environmental fluctuation magnitudes, and tallied the survivors (Fig 4A-C, Methods). These simulations included “anomalous” species (who resource preference orders did not match the ordering of their growth rates) in initial species pools, but, consistent with previous reports [57], these anomalous species almost never survived (Appendix §6), thus making them largely irrelevant. Without environmental fluctuations, all 100 communities saturated but never exceeded competitive exclusion (Fig 4C). However, with even small fluctuations of σ_RS_ = 0.09 (Fig 4B for illustration), 74/100 random communities violated the competitive exclusion principle (Fig 4C). For fluctuations between σ_RS_ = 0.09 and σ_RS_ = 0.236 diversity remained high, with 64/100 to 83/100 communities violating competitive exclusion and an average of 4.21 +/-0.04 coexisting survivors (on just the three resources). Thus, with even small to moderate magnitudes of environmental fluctuation, the majority of random communities violated the competitive exclusion principle.

**Figure 4.**
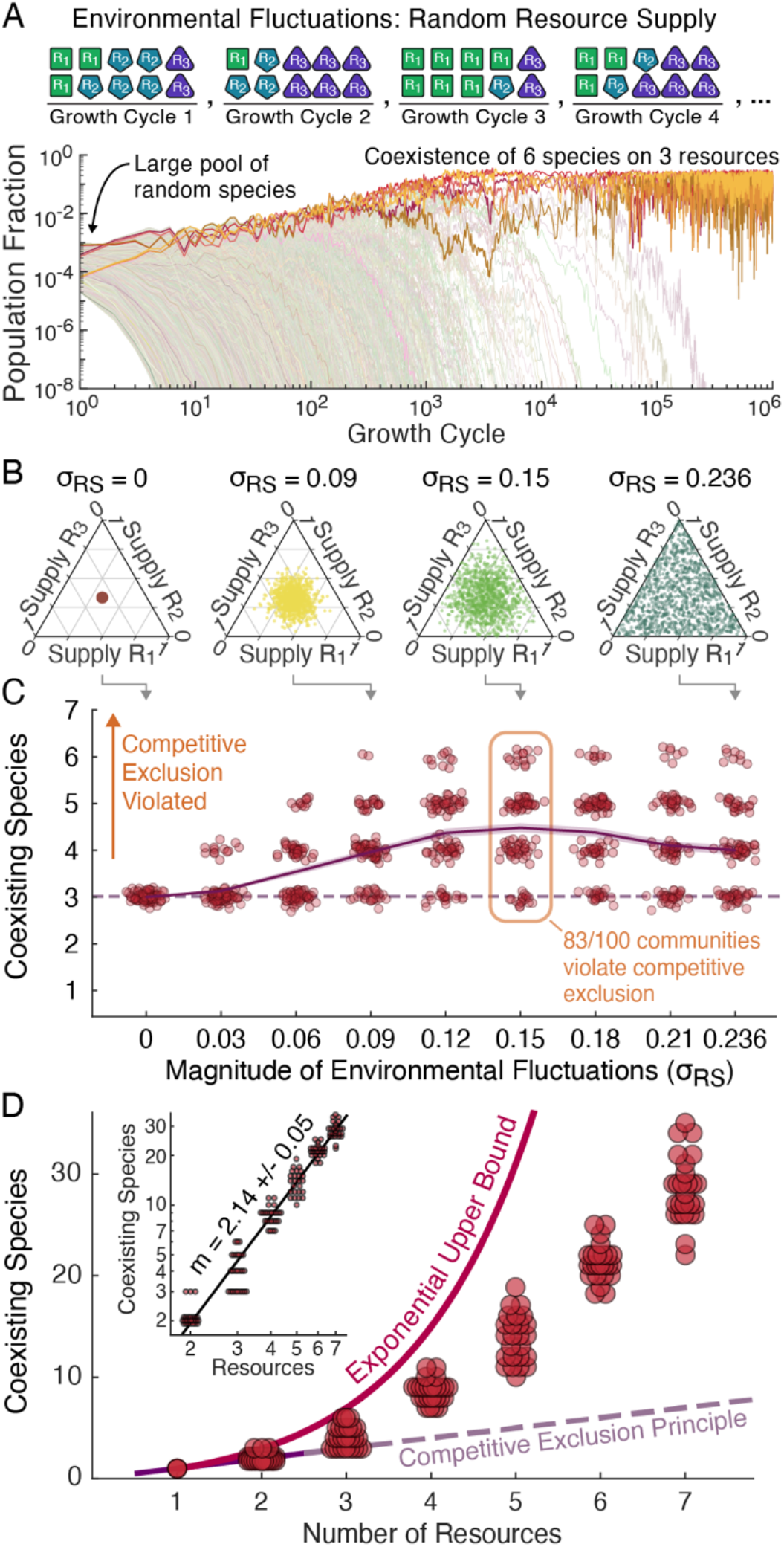
With environmental fluctuations, communities of random species frequently violate competitive exclusion. (**A**) Top row illustrates our implementation of extrinsic environmental fluctuations by randomly sampling the resource supply fractions on each growth cycle while keeping total supply constant (Methods). Plot shows population fractions at the end of each growth cycle for an example in which 5,000 random species were inoculated and simulated for 10^6^ growth cycles with a uniform-randomly sampled resource supply on each day (σ_RS_ = 0.236). (**B**) Illustration of our resource supply sampling distributions as a function of the fluctuation magnitude σ_RS_ on simplex diagrams in which an equal supply of all resources maps to the middle of the triangle and a single resource being supplied maps to a corner (Appendix §1.2, Methods). For σ_RS_ = 0.09, 0.15, and 0.236, one thousand randomly sampled resource supplies are shown. (**C**) We sampled 100 communities of 5000 species competing for three resources with the metabolic constraint that the L2-norm of species’ growth rates equal 1 hr^-1^, simulated these communities at 9 different fluctuation magnitudes, and tallied the survivors (Methods). Points represent the number of coexisting species for each community at each fluctuation magnitude, the dashed line shows the competitive exclusion principle (one species for each of the three resources), and the solid line and shaded region provide the mean number of survivors and standard error. With even the smallest level of fluctuation 13/100 of communities violated competitive exclusion, and with fluctuations of σ_RS_ = 0.09 or larger competitive exclusion was usually violated, with a peak frequency of violations of 83/100 at σ_S R_= 0.15. (**D**) We next simulated 25 random species pools for each of two through seven resources using a uniform-random resource supply (σ_RS_ = 0.236 for the three-resource case, Methods). Points represent the number of survivors in the simulations. Shown for comparison are the competitive exclusion principle (one species per resource) and the exponential upper bound (Fig 3B). Inset shows the same data on a log-log scale with a monomial best fit. The monomial was a better fit than both linear and exponential models (Appendix §1.3).

Having shown competitive exclusion violations to be the most likely outcome with a fluctuating three-resource supply, we also investigated community diversity with more than three resources available. We sampled random growth rates for large species pools growing on two through seven resources (Methods) and simulated these communities with a uniform-random resource supply (σ_RS_ = 0.236 in the three-resource case). As anomalous species had never survived under a uniform-random three-resource supply in previous simulations (Appendix §6) they were not included in the species pools for this next batch of simulations. With four or more resources, all random communities violated the competitive exclusion principle with up to several times as many species as resources coexisting (Fig 4D). For example, with seven resources we saw an average of 28.2 +/-0.6 survivors or four stably coexisting species for every resource (Fig 4B). However, while the expected number of survivors increased rapidly with the number of resources, it did not grow exponentially as our previously derived upper bound suggested it might. Instead, the best fit through our data grew slightly faster than quadratically, specifically 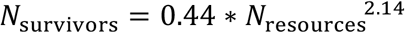 (Fig 4D). This growth in the number of coexisting species nevertheless scales to explain massively diverse communities, for example predicting over 100 species should stably coexist on 13 resources and over 1000 on 38. All random communities violating competitive exclusion by considerable margins indicates that resource competition can be sufficient to explain highly diverse communities.

### Temporal niches explain how environmental fluctuations reshape community composition

With highly diverse communities shown to be a likely outcome of diauxic resource competition, we proceeded to explore whether our understanding of temporal niches could be used to predict the survivors and structure of these communities. To do this, we returned to the 100 random communities of 5000 species competing for 3 resources (Fig 4C), extended our definition of environmental fluctuations to allow for magnitudes up to σ_RS_ = 0.471, which corresponded to a single randomly selected resource being supplied each growth cycle (Fig 5A, Methods), and simulated the communities at eight additional fluctuation magnitudes from σ_RS_ = 0.27 to σ_RS_ = 0.471.

**Figure 5.**
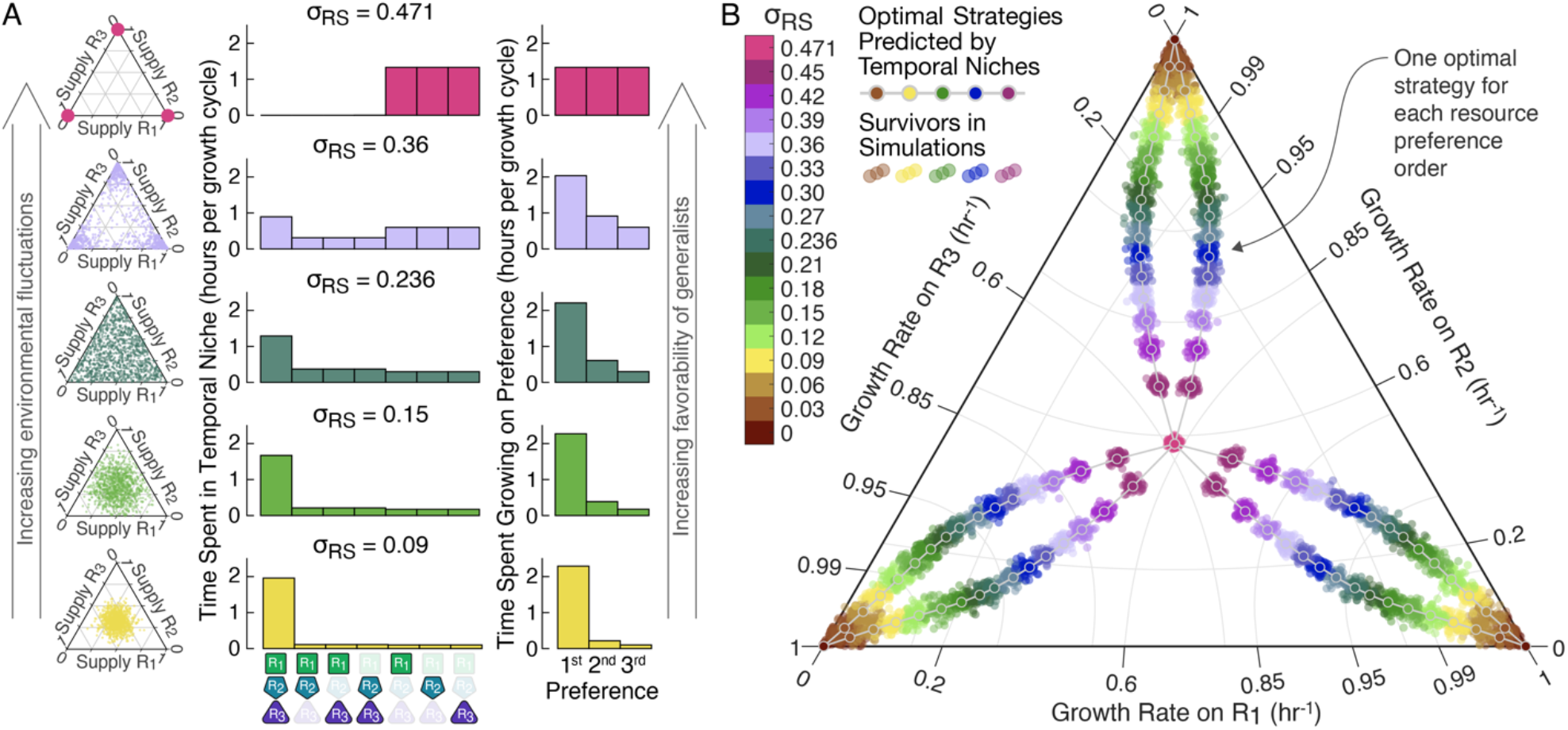
Temporal niches explain the composition of highly diverse communities in fluctuating environments. (**A**) We extended the resource supply sampling distributions to now have a maximum value σ_RS_ = 0.471, corresponding to a single randomly supplied resource on each growth cycle. Resource supply distributions are shown in the left column on the same simplex diagrams as in Figure 4B. We then simulated the 100 random communities of 5000 species from Figure 4 at 8 additional magnitudes for 17 total fluctuation magnitudes from σ_RS_ = 0 to σ_RS_ = 0.471. To develop a prediction of optimal strategies, we began by looking at the time spent in each temporal niche (middle column) and the time species spent growing on each of their resource preferences (right column). As the environmental fluctuation magnitude increased so did the time species spent growing on their second and third preferences, which would favor resource generalists. (**B**) Because species had been sampled with a metabolic constraint on their growth rates 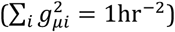, we could predict the optimal strategies (Appendix §2.4). We compared these optimal strategies against the survivors in simulations using a simplex plot which is nonlinear in *g*_*μi*_ but in which complete specialization on a single resource still maps to the corners and equal investment in all resources still maps to the center (Methods, Appendix §1.2). At all fluctuation magnitudes, survivor growth rates are tightly clustered around the predicted optimal strategies.

We then considered the average time spent growing in each temporal niche and average time species spent growing on each preference (Fig 5A). Without fluctuations, all resources were depleted at nearly the same time, as is consistent with previous reports [57]. This nearly simultaneous depletion meant essentially all growth occurred in the all-resources niche with all species growing on their top preferences, favoring specialists entirely invested in their top preference. By contrast, with only a single random resource supplied on each growth cycle at the maximum magnitude of environmental fluctuation, species spent equal time growing on each resource, favoring generalists equally invested in all resources (Fig 5A, top row). At intermediate levels of environmental fluctuations, the fluctuating depletion times meant all temporal niches occurred and that species grew on each resource but with more time spent on top preferences, favoring intermediate strategies in which species had some investment in all resources but greater investment on higher preferences (Fig 5A middle and bottom rows).

We next calculated the optimal allocation of a species’ growth rate allowance amongst its first, second, and third preferences at each fluctuation magnitude (Fig 5B, Methods) based on the expected time on each preference and the metabolic constraint species had been sampled with 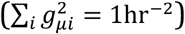. This optimal allocation could be applied to each of the possible resource preference orders to predict a set of six optimal strategies that would together form an uninvadable “supersaturated” community (Fig 5B, Appendix §2.4). We then compared the predicted optimal strategies to the growth rates of the species that had survived in simulation. At all fluctuation magnitudes the observed survivors had growth rates tightly clustered around the optimal strategies (Fig 5B). Our understanding of temporal niches thus led to intuitive and accurate predictions of the survivors in highly diverse communities, highlighting the potential for temporal niches as a lens for understanding the competitive structure of highly diverse communities.

### Temporal niches predict diversity under periodic oscillations

We concluded our study by considering community assembly under periodic environmental fluctuations. We worked with the case of three resources and defined three environmental oscillations of different periods (Appendix Figure S13A, Methods). We also noted that within our model, death does not need to occur between each growth cycle but could instead occur once per environmental oscillation with the model yielding the exact same results. For example, the period-three oscillation could be viewed as growth on fresh pulses of resources in spring, summer, and fall seasons followed by a winter season in which substantial death occurs. Using our understanding of temporal niches, we predicted (i) high diversity under the period-three oscillation due to the supply changing significantly on each growth cycle, (ii) even higher diversity under the period-six oscillation due to the addition of more distinct resource supplies, but (iii) less diversity under the period-twelve oscillation due to the supply now changing only relatively little from one growth cycle to the next (Appendix §5). Consistent with these predictions, we observed an average of 4.5 survivors in the three-season oscillation, 5.3 survivors in the six-season oscillation, and 3.3 survivors in the twelve-season oscillation (Appendix §5, Appendix Figure S13B, Methods). Thus, the temporal niches that arise from sequential resource utilization provide insights into highly diverse communities formed under all three types of fluctuation considered: internally driven oscillations and chaos, stochastic environmental fluctuations, and seasonal or other periodic environmental oscillations.

## Discussion

In this paper, we used a model of sequential resource utilization to explore highly diverse communities first with community-driven oscillations under constant environmental conditions (Fig 1-3), then with environmental fluctuations in the form of a stochastic resource supply (Fig 4-5), and finally with a periodically varying resource supply (Appendix Fig S13). We used the case of community-driven oscillations to derive a temporal niche structure capable of supporting exponentially many species as resources (Fig 2-3) and showed that the same temporal niches explained community structure under all three types of fluctuation. We demonstrated that with environmental fluctuations most random communities competing for three resource and essentially all communities competing for four or more resources will violate the competitive exclusion principle with up to several times as many coexisting species as resources (Fig 4). The temporal niche structure derived from the case of community-driven oscillations accurately predicted which species would survive in the case of environmental fluctuations (Fig 5). We confirmed the importance of sequential utilization specifically by demonstrating that a model with sequential utilization removed and all other dynamics left as intact as possible cannot support more species than resources (Appendix §7). Our results thus demonstrate that sequential resource utilization and environmental fluctuations robustly produce highly diverse communities and that the structure of these communities can be accurately predicted by temporal niches.

While we considered the simplest model of diauxic growth, numerous variations could be explored. For example, we did not incorporate diauxic lags, which are periods of little to no growth as species switch resources and which have been shown to be a source of coexistence [49,59,68,69]. If a species sometimes but not always finishes its lag in time to grow in a specific temporal niche, this sometimes-missed niche would be effectively split in two, potentially greatly increasing expected and maximum diversity. Similarly, Monod growth dynamics, which have been shown to support multiple coexisting species on a single resource [70], might increase diversity in our model by having some species perform best early in a temporal niche while others perform best later. Additionally, microbial growth typically displays a mix of sequential and simultaneous utilization, consuming multiple but not all resources at a time [53,56,59]. If species’ simultaneous utilization granted independent growth rates in each niche then communities that saturate the upper bound on diversity could become a common occurrence due to potentially having a different species be the fastest grower in each niche. However, the temporal niche structure we have developed in this paper would continue to accurately describe community dynamics. Finally, some resources could be cross-fed such that niches containing the cross-fed resources would emerge only after degradation of the supplied resources. This cross-feeding could add variety to the order in which temporal niches occur and even allow for more niches than resources to be realized on a single growth cycle (for example only the supplied resource then both resources then only the cross-fed resource). Incorporating each of these growth phenomena into the diauxie model could yield even greater diversity, but with temporal niches remaining a powerful framework for understanding community dynamics.

The metabolic constraint with which we sampled species could also be modified or even eliminated. When considering simultaneous utilization of multiple resources, metabolic constraints are an intuitive property of microbes as producing the enzymes to consume one resource leads to less capacity to produce the enzymes for other resources [44,50,56,71–73]. With diauxie, however, microbes invest entirely in one resource at a time, so the origin of a tradeoff becomes unclear, although it still seems unlikely to have best-at-everything competitors. Concluding that best-at-everything species should not occur but investment in one resource should not necessarily take away from investment in another in a one-to-one fashion motivated us to select an L2-norm for sampling species rather than the L1-norm typically used for tradeoffs in simultaneous utilization. Macroscopic organisms, by contrast, cannot rapidly retool to the same extent microbes can and would therefore more likely be constrained by competitive tradeoffs. Nevertheless, to confirm having a metabolic constraint wasn’t a prerequisite for our results, as is the case for other demonstrations of highly diverse communities [23,37,41,43,44,74,75], we ran additional simulations without any sampling constraints and saw that competitive exclusion violations were still a likely outcome (Appendix §4.1). These simulations thus confirm that highly diverse communities enabled by temporal niches are a likely outcome even without metabolic constraints.

Similarly, in nature all resources are not equally abundant, nutritious, or likely to be preferred over each other [60], motivating an investigation into the robustness of our results to these asymmetries. To test robustness against asymmetries in resource abundance and nutritional value, we ran simulations in which some resources were supplied in greater average amounts and in which some resources supported faster growth rates and again saw competitive exclusion violations remain likely (Appendix §4.2). To explore the effect of some preference orders being more likely than others, we leveraged recent results from Takano et al. on how the structure of central metabolism makes some resource preference orders much more likely to appear than others [60]. Specifically, for each set of three of the seven resources Takano 2023 focuses on, we used the relative frequencies of the most common seven-resource preference orders (Fig 1E in Takano 2023) to determine the relative frequencies of the three-resource preference orders, sampled species from that distribution, and ran additional simulations (Appendix §7). For some selections of three resources only a single preference order was possible and diversity was low with sometimes only a single species surviving, but for other selections of resources as many as five of the six preference orders were possible and up to six species coexisted on the three resources (Appendix §7). Thus, our results appear generally robust to various possible resource asymmetries.

Final communities in the fluctuating environments featured species that naturally clustered due to the assumption of sequential resource utilization. Specifically, optimal strategies were to invest primarily in one’s first preference and by an amount that does not depend on the order of later preferences (and in general to invest in the first *N* preferences by amounts that do not depend on later preferences). Thus, species all sharing the same first preference form a natural cluster. This clustering reflects “lumpy coexistence”, the phenomenon of communities displaying multiple “lumps” or clusters of similar species interacting nearly neutrally within a lump but with strong inter-lump competition [37,40,42,76–78]. For communities growing on several resources an interesting structure of lumps-within-lumps emerges with the coarsest lumps being the species all sharing the same first preference and the increasingly fine-scale lumps being the species all sharing their first *N* preferences. Consistent with strong inter-lump but weak intra-lump competition, we observed all simulated communities to have at least one species with each resource as its top preference but a decreasing complementarity of *N*^th^ preferences amongst species that shared their first *N*–1 preferences (Appendix §6). This pattern of lumps-within-lumps could be an interesting avenue of future research and should be readily testable in experimental data.

Temporal niches supporting increased diversity in a fluctuating environment can be understood as a mechanistic basis for realizing the storage effect [18,25,79–81] on the scale of many species and resources. The storage effect is a statistical bias towards coexistence when species have temporally fluctuating fitness, persistence through unfavorable periods, and nonlinearities that create increasing potential for large rebounds in populations size as their relative abundance decreases. It has been thoroughly explored in the case of a couple species coexisting due to fluctuations in one or a couple environmental variables, but a noteworthy question remains how a large number of species could each have the potential for sufficiently beneficial conditions to survive [82]. In communities of sequential utilizers each following distinct optimal strategies, a species will be the fittest of all its competitors when the relative supply of each resource matches its resource preference order (i.e. its top preference is supplied in the greatest amount, its second in the second greatest amount, etc.). Because there are many more optimal strategies then available resources, our modeling explains how a massive number of competing species could each have the potential for sufficiently beneficial conditions on particular growth cycles to recover and avoid extinction. Ecologists should therefore consider whether a species that appears to lack the fitness necessary to coexist with its competitors benefits when resources are available in specific relative abundances and specific temporal niches are created.

Our findings about temporal niches suggest several phenomena and properties which should be relevant across natural ecosystems at both microbial and macroscopic scales. The fundamental elements of diauxie are present whenever an ecosystem experiences boom and bust cycles or periodically available resources. For example, gut microbes breaking down food when their hosts have a meal, algae blooming after significant rainwater runoff, and foragers feasting when foliage blooms all experience sudden resource influxes and eventual depletions. In all cases, some resources will inevitably be depleted before others and a temporal niche structure will be created. Variability in relative resource influxes readily creates variability in relative resource depletion rates, which makes not just how a species responds to a depleting environment but how its response depends on which resources are being depleted most rapidly an important and consequential property. As we have shown in this paper, small changes in initial resource availability can create fundamentally different temporal niches as the environment becomes depleted. Our work highlights how these temporal niches can explain highly diverse communities in a wide range of ecosystems with many different forms of fluctuation. We hope this work can be a useful guide for other researchers exploring highly diverse communities, particularly those that appear to be driven by the competition for a much smaller number of resources than coexisting species.

## Methods

### Diauxie Model

For clarity, we have used Greek letters (*μ, ν*, etc.) whenever indexing over species and Latin letters (*i, j*, etc.) whenever indexing over resources. Units are dimensionless except for time.

Species have population size *n*_*μ*_(*t*) and growth rates *g*_*μi*_. Resources have concentrations *c*_*i*_(*t*). Species grow exponentially on whichever resource is their highest remaining preference and consume one dimensionless unit of resource for each dimensionless unit of biomass gained (i.e. all yields equal 1). This produces dynamical equations:

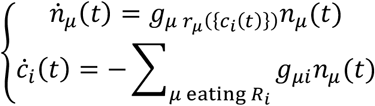

where the notation *r*_*μ*_({*c*_*i*_(*t*)}) indicates the index of the resource species *μ* is eating given the current resource concentrations (i.e. which resource ins its top remaining preference).

Growth rates and population sizes are always positive, so ċ_*i*_ (*t*) ≤ 0 and eventually *c*_*i*_(*t*) = 0 is reached for each resource. We label the time when *c*_*i*_(*t*) = 0 is reached for a particular resource as its depletion time *t*_dep*i*_. When all resources have been depleted, the resource concentrations are reset to *c*_*i*_ → *s*_*i*_ where {*s*_*i*_} are the supply concentrations and all population sizes experience a mortality 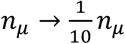. We refer to the time between consecutive resets as one growth cycle.

In all our simulations ∑_*i*_ *s*_*i*_ = 1, so at the end of a growth cycle 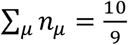 after accounting for day-to-day carryover. Reported population fractions are always at the end of growth cycle and therefore equal to 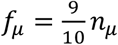.

### Numerical Simulations

Simulations were performed in MATLAB. Because all growth is exponential, calculation of resource depletion times and species’ growth on each cycle could be achieved by solving a sum of exponentials equations, eliminating the need for integrating dynamical equations. Species were declared extinct and removed from simulations when their population fractions dropped below 10^-12^. Code needed to reproduce the basic results of the paper will be made available at the time of final publication.

### Sampling Species in Environmental Fluctuation Simulations

Species were uniform-randomly sampled from the surface defined by 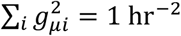 (where *i* indexes over a species’ growth rates on different resources) by sampling from a symmetric multi-dimensional normal distribution, taking the absolute value of each component, and normalizing growth rate vector lengths to 1 hr^−1^. (This L2 norm (rather than the L1 commonly used in simultaneous utilization models) disallowed best-at-everything species while capturing that, because species are not consuming resources at the same time, investment in one resource should not necessarily take away from the ability to invest in another at a later time in a one-to-one fashion.)For the simulations with three resources and a varying fluctuation magnitude, 5000 species were sampled with independent growth rates and resource preferences, allowing so-called “anomalous” species. The inclusion of these anomalous species was, however, largely inconsequential as for most conditions no anomalous species survived (Appendix §6). The same 100 pools of 5000 species per pool were used at all fluctuation magnitudes.

For the simulations with a variable number of resources, growth rates were randomly sampled and resource preferences were then set as matching the each species’ growth rate order (thus disallowing “anomalous” species, which had been exceedingly rare in the previous set of simulations). To adjust for sampling species from a larger parameter space as the number of resources was increased, we held 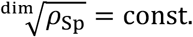, where dim = *N*_Re_ − 1 was the dimension of the space species were being sampled from and 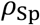 was the density of species, and adjusted the number of species accordingly. The constant value of 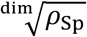 was set such that the three-resource case used pools of 500 species.

### Fluctuating Resource Supply Probability Distributions

In simulations in which the resource supply was not fluctuating, all resources were supplied in equal amounts.

When there were environmental fluctuations, resource supplies on each growth cycle were sampled from maximum entropy distributions under the constraint that on each growth cycle ∑_*i*_ *s*_*i*_ = 1 where *s*_*i*_ is the supply of resource R_*i*_ and that averaged across growth cycles the fluctuation magnitude,

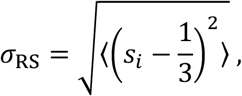

had the desired value. These sampling distributions had the functional form of a normal distribution that was centered on an equal supply of all resources, was clipped by the boundary *s*_*i*_ > 0, and gained a negative variance at large fluctuation magnitudes. At small values the environmental fluctuation magnitude and the variance of the normal distribution were equal, but as clipping by the boundary *s*_*i*_ > 0 became significant the variance of the normal distribution grew faster than the fluctuation magnitude.

For the analysis of how diversity increased with increasing fluctuation magnitude (Fig 4), values of σ_RS_ > 0.236 were excluded. The value 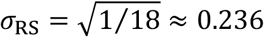 corresponds to a uniform-random sampling of {*s*_*i*_} so within the region 0.236 < σ_RS_ ≤ 0.471 the entropy of the sampling distribution declines with increasing σ_RS_. (For example, the limiting case of a single resource on each day at σ_RS_ = 0.471 is arguably a less random resource supply than the uniform-random sampling at σ_RS_ = 0.236 despite having a larger fluctuation magnitude as defined.) This unnecessarily complicated the interpretation of σ_RS_ > 0.236, so this region was excluded from the analysis of how diversity increased with increasing fluctuation magnitude (Fig 4C). The full range 0 ≤ σ_RS_ ≤ 0.471 was, however, used for the analysis of how optimal strategies changed with increasing fluctuation magnitude (Fig 5) in order to highlight how an absolute generalist strategy eventually becomes favorable at σ_RS_ = 0.471. For the simulations involving a variable number of resources (Fig 4D), the resource supply was always sampled uniform-randomly.

### Numerically Sampling Resource Supply Distributions

For three resources, numerical sampling was implemented at small fluctuation magnitudes by sampling from normal distributions (using a lookup table to determine the correct variance after accounting for clipping by *s*_*i*_ > 0) and rejecting and resampling values outside *s*_*i*_ > 0. For three resource and intermediate to large fluctuation magnitudes (σ_RS_ ≥ 0.21), it proved faster to use a rejection-based sampling in which we uniform-randomly sampled *s*_1_ through *s*_2_, calculated *s*_3_ = 1 − *s*_1_ – *s*_2_, rejected and resampled if *s*_3_ < 0, and then randomly rejected and resampled at probability

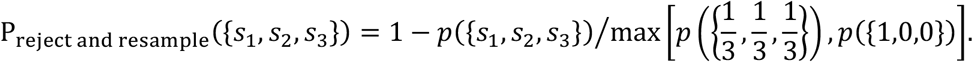

### Repetition of Simulations with Fluctuating Resource Supplies

For the simulations with three resources and a varying fluctuation magnitude, simulations were repeated five times in case species that could have coexisted with the other survivors experienced early stochastic extinctions. In many cases it appeared that one species would survive only if another experienced an early extinction. To determine which of these species was fitter, we then repeated simulations another three times, now giving any species that had survived in any of the first five simulations a head start (with population sizes ∼10^3^x higher) over the species that had been consistently excluded. This also served to test whether the final communities were vulnerable to invasion. The reported survivors are those that survived in at least two out of the last three repetition of the simulations. Time traces were then visually inspected to remove any species that were clearly trending towards but not quite reaching the extinction threshold (a population fraction of 10^-12^).

### Simplex Diagrams

For a visual explanation of simplex diagrams (used in Figures 4B and 5) see Appendix §1.2 and Appendix Figure S1.

### Best Fit for Number of Species vs Number of Resources

We test two-parameter linear, monomial, and exponential fits. The monomial fit was the best fit as measured by either R^2^ or mean square error:

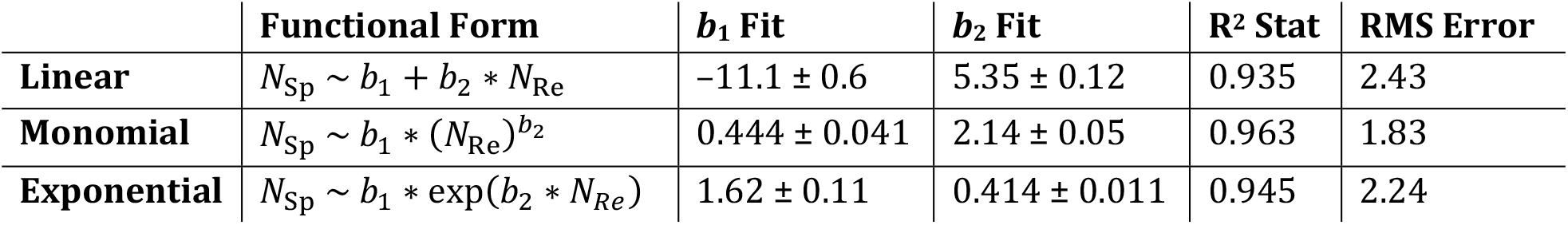

However, the monomial being the better fit was not statistically significant as measured by a F statistic: *p* ≈ 0.36 for monomial vs linear and *p* ≈ 0.40 for monomial vs exponential.

### Down-Sampling in Figure 4A Plot

Due to computer memory limitations, we could not plot the full trajectories (including all 10^6^ timesteps for all 5000 species), so for generating the plot shown in Figure 4A data were down-sampled. Population fractions after each of the first ten growth cycles are all shown. Populations fractions are then shown for every 2^nd^ growth cycle for cycles 10 through 50. Then every 5^th^ cycle for cycles 50 through 200. Then every 10^th^ for cycles 200 through 1000. Every 20^th^ for 1,000 through 2,000. Every 50^th^ for 2,000 through 4,000. Every 100^th^ for 4,000 through 7,000. Every 200^th^ for 7,000 through 10,000. And every 500^th^ for 10,000 through 1,000,000.

### Periodic Environmental Oscillations

The periodic (aka seasonal) resource supplies were calculated from sinusoidal oscillations, which also traced circles on the simplex plots used (Appendix Fig S13A). The specific resource supplies were:

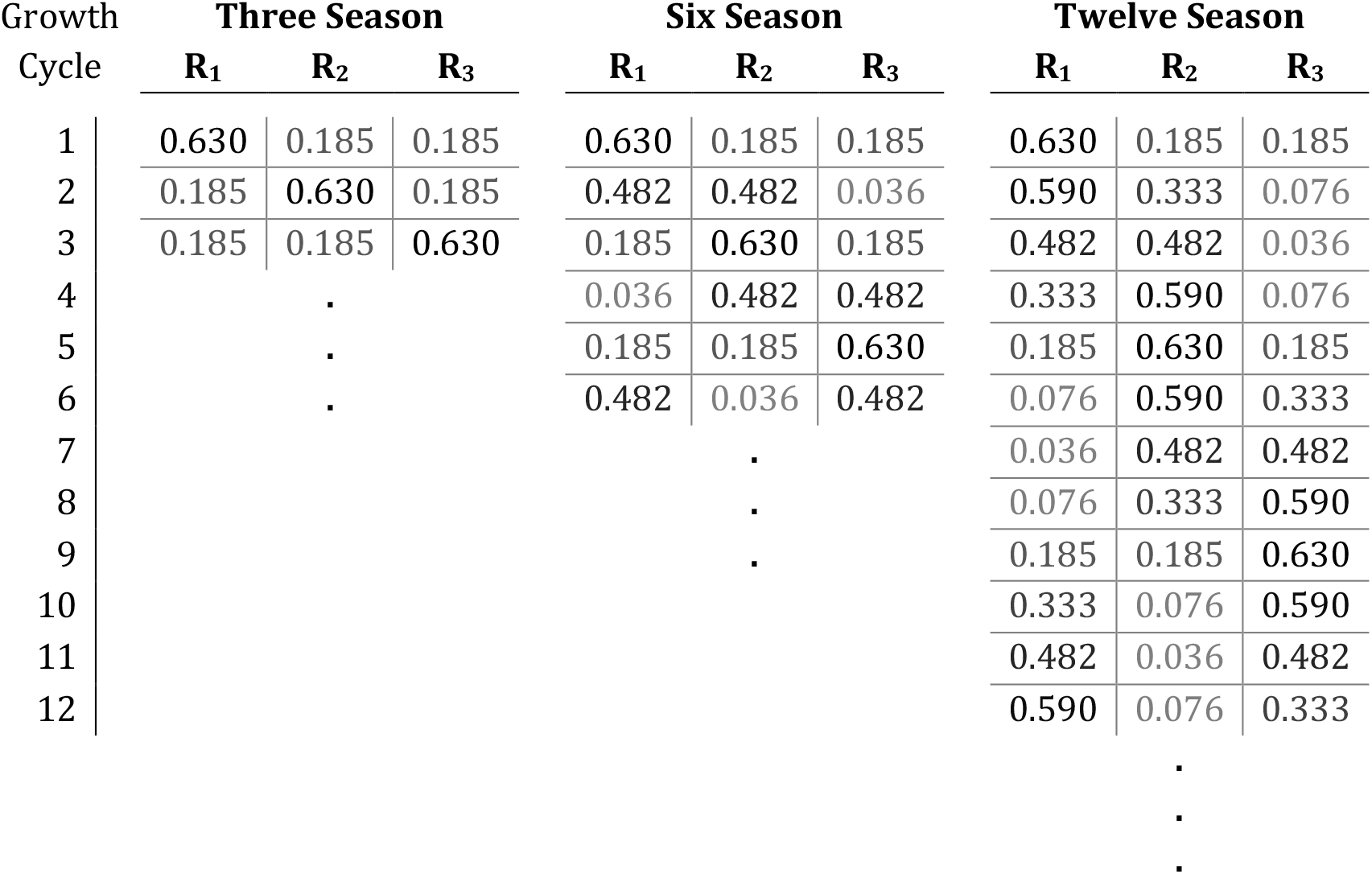

| Growth<br>Cycle | Three Season |  |  | Six Season |  |  | Twelve Season |  |  |
| --- | --- | --- | --- | --- | --- | --- | --- | --- | --- |
|  | R<sub>1</sub> | R<sub>2</sub> | R<sub>3</sub> | R<sub>1</sub> | R<sub>2</sub> | R<sub>3</sub> | R<sub>1</sub> | R<sub>2</sub> | R<sub>3</sub> |
| 1 | 0.630 | 0.185 | 0.185 | 0.630 | 0.185 | 0.185 | 0.630 | 0.185 | 0.185 |
| 2 | 0.185 | 0.630 | 0.185 | 0.482 | 0.482 | 0.036 | 0.590 | 0.333 | 0.076 |
| 3 | 0.185 | 0.185 | 0.630 | 0.185 | 0.630 | 0.185 | 0.482 | 0.482 | 0.036 |
| 4 |  | . |  | 0.036 | 0.482 | 0.482 | 0.333 | 0.590 | 0.076 |
| 5 |  | . |  | 0.185 | 0.185 | 0.630 | 0.185 | 0.630 | 0.185 |
| 6 |  | . |  | 0.482 | 0.036 | 0.482 | 0.076 | 0.590 | 0.333 |
| 7 |  |  |  |  | . |  | 0.036 | 0.482 | 0.482 |
| 8 |  |  |  |  | . |  | 0.076 | 0.333 | 0.590 |
| 9 |  |  |  |  | . |  | 0.185 | 0.185 | 0.630 |
| 10 |  |  |  |  |  |  | 0.333 | 0.076 | 0.590 |
| 11 |  |  |  |  |  |  | 0.482 | 0.036 | 0.482 |
| 12 |  |  |  |  |  |  | 0.590 | 0.076 | 0.333 |
|  |  |  |  |  |  |  | . |  |  |
|  |  |  |  |  |  |  | . |  |  |
|  |  |  |  |  |  |  | . |  |  |

## Acknowledgements

We thank members of the Gore Lab and Simons Foundation Principles of Microbial Ecology collaboration for discussion.

## Author Contributions

Conceptualization: BB, HL, JG

Methodology: BB, HL, JG

Software: BB

Investigation: BB

Formal analysis: BB

Visualization: BB

Writing–original draft: BB, HL, JG

Writing–review and editing: BB, HL, JG

## Data Availability

All data are available in the main text or supplementary materials along with detailed descriptions of all simulations and parameters choices for reproducing all specific examples.

