## Appendix for "Biodiversity is enhanced by sequential resource utilization and environmental fluctuations via emergent temporal niches"

*for*


#### §1 Additional Information for Main Text figures

- §1.1 Simulation parameters for Main Text figures.
- §1.2 Simplex Plots
- §1.3 Linear vs monomial vs exponential fits in Figure 4D
- §1.4 Number of surviving species including data at larger fluctuation magnitudes

#### §2 Properties of the Diauxie Model

- §2.1 Detailed definition of the model and importance of resource depletion times
- §2.2 At a fixed point, competitive exclusion is obeyed
- §2.3 With fluctuations, exponentially many species as resources can coexist
- §2.4 Optimal strategies

#### §3 Additional Results with a Constant Resource Supply

- §3.1 With three species competing for two resources a fixed point was always reached
- §3.2 When sampling random communities, emergent oscillations are rare
- §3.3 Example of five species coexisting on four resources without anomalous resource preferences
- §3.4 Example of chaos
- §3.5 Examples of seven species coexisting on three resources, fifteen coexisting on four, and twenty-three on five
- §3.6 Intuitive construction of an oscillating community
- §3.7 Robustness against demographic noise

#### §4 Additional Results with a Variable Resource Supply

- §4.1 Competitive exclusion violations are a likely outcome even without metabolic constraints
- §4.2 Competitive exclusion violations are a likely outcome even if some resources support universally faster growth rates than others
- §4.3 Sampling from evidence-based resource preference order distributions

#### §5 Seasonal Environmental Cycles

#### §6 Resource Preference Complementarity and Anomalous Species

#### §7 Comparison to a Model Without Sequential Resource Utilization

### §1 Additional Information for Main Text figures

#### §1.1 Simulation parameters for Main Text figures.

In Figures 1 and 2, resources are supplied in equal amounts with $\sum_i s_i = 1$ at the start of each day and the dilution factor is $\mathrm{DF} = 10$. In Figure 1A, the two species are started from equal population sizes $n_{\mathrm{A}}^{\mathrm{day}\ 1}(0) = n_{\mathrm{B}}^{\mathrm{day}\ 1}(0) = \frac{1}{2(\mathrm{DF}-1)}$ and have growth rates and resource preferences as shown in the figure:

| | $R_1$ | $R_2$ |
|---|---|---|
| Species A | 0.75 $\mathrm{hr}^{-1}$ | 0.1 $\mathrm{hr}^{-1}$ |
| Species B | 0.15 $\mathrm{hr}^{-1}$ | 0.45 $\mathrm{hr}^{-1}$ |

**Table S1.** Figure 1A growth rates.

| | 1st Pref | 2nd Pref |
|---|---|---|
| Species A | $R_1$ | $R_2$ |
| Species B | $R_2$ | $R_1$ |

**Table S2.** Figure 1A preferences.

In Figure 1B, the five species are start from equal population sizes $n_{\mu}^{\mathrm{day}\ 1}(0) = \frac{1}{5(\mathrm{DF}-1)}$ in both the single-resource and two-resource examples. Growth rates in the one-species example are:

| | |
|---|---|
| Species A | 1.00 $\mathrm{hr}^{-1}$ |
| Species B | 0.85 $\mathrm{hr}^{-1}$ |
| Species C | 0.75 $\mathrm{hr}^{-1}$ |
| Species D | 0.60 $\mathrm{hr}^{-1}$ |
| Species E | 0.40 $\mathrm{hr}^{-1}$ |

**Table S3.** Figure 1B single resource example growth rates.

Growth rates and resource preferences in the two-species example are:

| | $R_1$ | $R_2$ |
|---|---|---|
| Species A | 1.00 $\mathrm{hr}^{-1}$ | 0.50 $\mathrm{hr}^{-1}$ |
| Species B | 0.60 $\mathrm{hr}^{-1}$ | 0.90 $\mathrm{hr}^{-1}$ |
| Species C | 0.85 $\mathrm{hr}^{-1}$ | 0.45 $\mathrm{hr}^{-1}$ |
| Species D | 0.10 $\mathrm{hr}^{-1}$ | 0.75 $\mathrm{hr}^{-1}$ |
| Species E | 0.60 $\mathrm{hr}^{-1}$ | 0.60 $\mathrm{hr}^{-1}$ |

**Table S4.** Figure 1B two-resource example growth rates.

| | 1st Pref | 2nd Pref |
|---|---|---|
| Species A | $R_1$ | $R_2$ |
| Species B | $R_2$ | $R_1$ |
| Species C | $R_1$ | $R_2$ |
| Species D | $R_2$ | $R_1$ |
| Species E | $R_2$ | $R_1$ |

**Table S5.** Figure 1B two-resource example resource preferences.

The example of five species surviving on three resources is the same in Figure 1C and Figure 2. The total population size is initially $\sum_{\mu} n_{\mu}^{\mathrm{day}\ 1}(0) = \frac{\mathrm{DF}}{\mathrm{DF}-1}$, and species' initial population fraction are:

| | |
|---|---|
| Species A | 0.65 |
| Species B | 0.19 |
| Species C | 0.08 |
| Species D | 0.03 |
| Species E | 0.05 |

**Table S6.** Figures 1C and 2 initial population fractions.

In the Figure 1C and 2 example, growth rates and resource preferences are:

| | $R_1$ | $R_2$ | $R_3$ |
|---|---|---|---|
| Sp. A | 0.452 | 0.829 | 0.195 |
| Sp. B | 0.315 | 0.465 | 0.855 |
| Sp. C | 0.464 | 0.402 | 0.477 |
| Sp. D | 0.484 | 0.463 | 0.145 |
| Sp. E | 0.250 | 0.433 | 0.478 |

**Table S7.** Figures 1C and 2 growth rates ($hr^{-1}$).

| | 1st Pref | 2nd Pref | 3rd Pref |
|---|---|---|---|
| Sp. A | $R_1$ | $R_2$ | $R_3$ |
| Sp. B | $R_2$ | $R_1$ | $R_3$ |
| Sp. C | $R_3$ | $R_1$ | $R_2$ |
| Sp. D | $R_1$ | $R_2$ | $R_3$ |
| Sp. E | $R_3$ | $R_2$ | $R_1$ |

**Table S8.** Figures 1C and 2 resource preferences.

In the Figure 1C and 2 example, species B and E never grow on $R_1$ in the steady-state oscillation, so their $R_1$ growth rates are irrelevant throughout Figure 2 (although they do grow on $R_1$ during the first couple days in Figure 1C).

In Figure 4A, all species were assigned random resource preference orders and growth rates uniform-randomly sampled from the surface defined by $\sum_i g_{\mu i}{}^2 = 1\ \text{hr}^{-2}$. The growth rates and resource preferences for the highlighted species (A, B, C, D, E, and F) are:

| | $R_1$ | $R_2$ | $R_3$ |
|---|---|---|---|
| Sp. A | 0.1250 | 0.2844 | 0.9505 |
| Sp. B | 0.2866 | 0.9483 | 0.1363 |
| Sp. C | 0.2582 | 0.1376 | 0.9535 |
| Sp. D | 0.0788 | 0.9608 | 0.2657 |
| Sp. E | 0.9460 | 0.1624 | 0.2657 |
| Sp. F | 0.9604 | 0.2540 | 0.1147 |

**Table S9.** Figure 4A growth rates (rounded to nearest 0.0001 $hr^{-1}$).

| | 1st Pref | 2nd Pref | 3rd Pref |
|---|---|---|---|
| Sp. A | $R_3$ | $R_2$ | $R_1$ |
| Sp. B | $R_2$ | $R_1$ | $R_3$ |
| Sp. C | $R_3$ | $R_1$ | $R_2$ |
| Sp. D | $R_2$ | $R_3$ | $R_1$ |
| Sp. E | $R_1$ | $R_3$ | $R_2$ |
| Sp. F | $R_1$ | $R_2$ | $R_3$ |

**Table S10.** Figures 4A resource preferences.

#### §1.2 Simplex Plots

Appendix Figure S1 explains the resource supply simplex plots presented in Main Text Figures 4B and 5A and Appendix Figure S4. Appendix Figure S2 explains the species growth rate simplex plot used in Main Text Figure 5B.

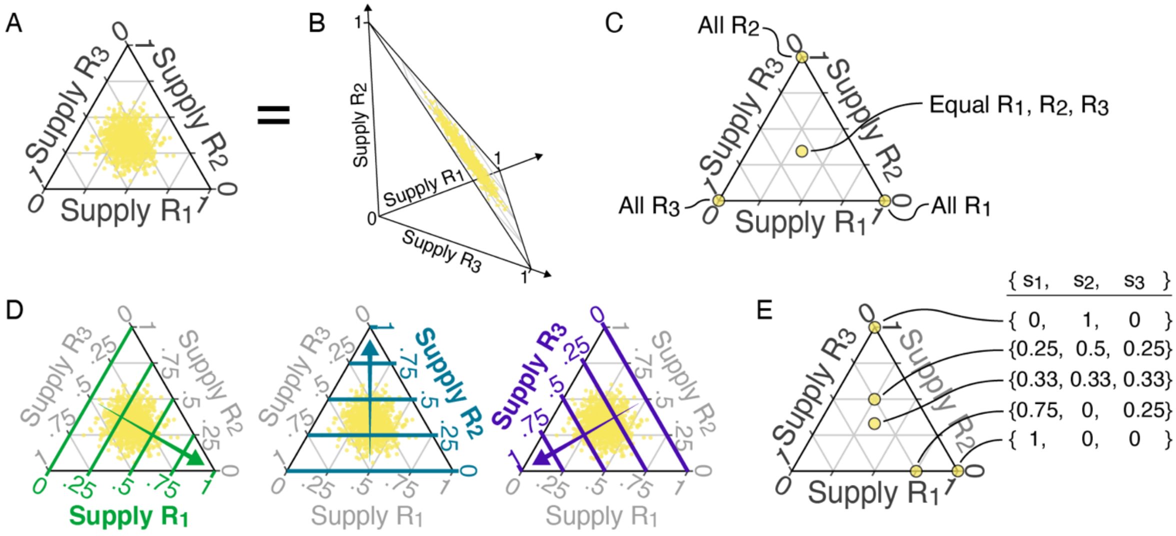


**Appendix Figure S1. Explanation of resource supply simplex plots.** (**A**) One of the simplex plots used to illustrate resource supply sampling distributions in the main text. (**B**) That simplex plot shown in three-dimensional $\{s_1, s_2, s_3\}$ space. The constraint $\sum_{i=1}^{\mathrm{N_{Re}}} s_i = 1$ means all resource supplies line on a two-dimensional surface. This surface is the simplex plot. (**C**) Qualitative relationships between position on the simplex plot and resource supply. The corners each correspond to the supply being entirely one resource. The center corresponds to the supply being equal amounts of each resource, the corners a single resource. (**D**) The simplex plot presented three times, each time highlighting a different component of the supply by bolding and coloring the grid lines corresponding to that resource's quantitative supply fractions. The supply of each resource increases linearly as the corner corresponding to the supply being entirely that resource is approached. (**E**) The quantitative supply fractions at five example points on the simplex.

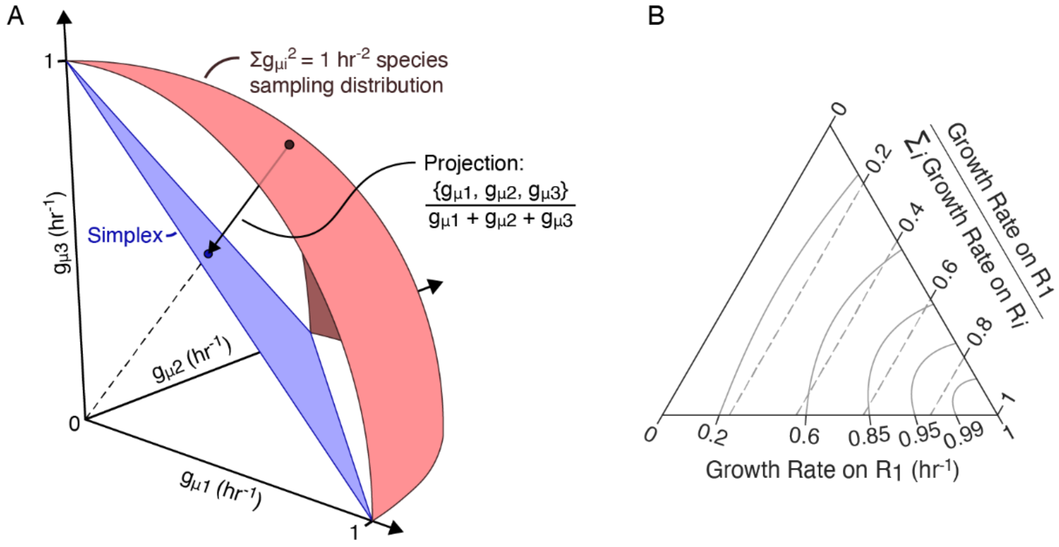


**Appendix Figure S2. Explanation of the growth rate simplex plots.** (**A**) The species sampling distribution in red and the simplex plot onto which growth rates were projected in blue. For the simulations with a fluctuating resource supply, species were sampled from $\sum_{i=1}^{N_{Re}} {g_{\mu i}}^2 = 1\ \mathrm{hr}^{-2}$. The nonlinearity of this constraint means species are not sampled from a flat surface (like the resource supply was) but instead from a curved surface, shown in red in $\{g_{\mu 1}, g_{\mu 2}, g_{\mu 3}\}$ space. This space is nevertheless two-dimensional (in the three-resource case), allowing species' growth rates to be projected onto a simplex plot with a bijective mapping. Various options for this projection exist. We choose to project growth rates allow vectors that pointed towards the origin $\{0, 0, 0\}$, as is illustrated by the arrow connecting a point on the growth rate sampling distribution (red) to the corresponding point on the simplex diagram (blue) and by the dashed line that continues this projection vector to the origin. (**B**) The nonlinearity of the $\sum_{i=1}^{N_{Re}} {g_{\mu i}}^2 = 1\ \mathrm{hr}^{-2}$ constraint and the projection used mean the simplex diagram is not linear in $\boldsymbol{g}_\mu = \{g_{\mu 1}, g_{\mu 2}, g_{\mu 3}\}$ but instead linear in $\boldsymbol{g}_\mu / \sum_{i=1}^{N_{Re}} g_{\mu i}$. The scale along the right edge of the simplex and the dashed lines across the simplex illustrate this linearity in $g_{\mu 1} / \sum_{i=1}^{N_{Re}} g_{\mu i}$. The scale along the bottom and curved lines across the simplex illustrate the corresponding nonlinearity in $g_{\mu 1}$.

#### §1.3 Linear vs Monomial vs Exponential Fits in Figure 4D

To test the scaling of the number of species with the number of resources, we fit linear, monomial, and exponential curves to the data. Appendix Table S11 summarizes the best fits, as well as $R^2$ and RMS residuals.

| | **Functional Form** | ***b*₁ Fit** | ***b*₂ Fit** | **R² Stat** | **RMS Error** |
|---|---|---|---|---|---|
| **Linear** | $N_{\mathrm{Sp}} \sim b_1 + b_2 * N_{\mathrm{Re}}$ | $-11.1 \pm 0.6$ | $5.35 \pm 0.12$ | 0.935 | 2.43 |
| **Monomial** | $N_{\mathrm{Sp}} \sim b_1 * (N_{\mathrm{Re}})^{b_2}$ | $0.444 \pm 0.041$ | $2.14 \pm 0.05$ | 0.963 | 1.83 |
| **Exponential** | $N_{\mathrm{Sp}} \sim b_1 * \exp(b_2 * N_{Re})$ | $1.62 \pm 0.11$ | $0.414 \pm 0.011$ | 0.945 | 2.24 |

**Appendix Table S11.** Details of the linear, monomial, and exponential fits to the data on number of surviving species vs number of resources presented in Main Text Figure 4D.

The ratio of mean square residual (MSR) of the monomial fit to the MSR of the linear fit is $\frac{1.83^2}{2.43^2} = 0.567$. This gives a F test p-value of p = 0.362. The ratio of MSR of the monomial fit to the MSR of the exponential fit is $\frac{1.83^2}{2.24^2} = 0.667$. This gives a F test p-value of p = 0.400. Thus, while the monomial fit is slightly better than the linear or exponential fits, it is not statistically significantly better. Appendix Figure S3 shows the linear, monomial, and exponential fits. Consistent with the statistical results, the monomial appears to be a visually slightly better fit, but the fits are all very similar across this range, so it is unsurprising that no fit is statistically significantly better.

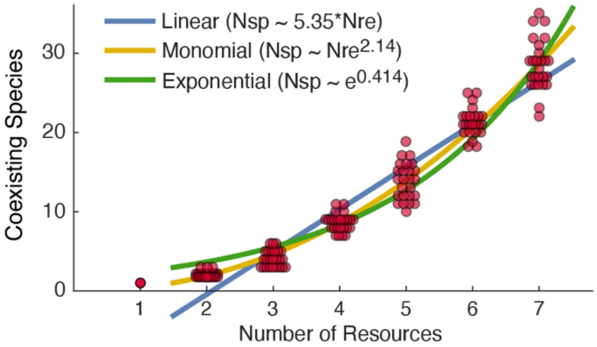


**Appendix Figure S3.** Linear, monomial, and exponential fits from Appendix Table S11 presented against the data from Main Text Figure 4D.

#### §1.4 Number of Surviving Species including Data at Larger Fluctuation Magnitudes

In Figure 4C (number of surviving species vs fluctuation magnitude), data for fluctuation magnitudes greater than $\sigma_{\mathrm{RS}} = 0.236$ was excluded because beyond this fluctuation magnitude the entropy of the sampling distribution actually decreases with increasing fluctuation magnitude. This decreasing entropy means it is ambiguous whether fluctuations should really be thought of as larger or the resource supply as more random beyond $\sigma_{\mathrm{RS}} = 0.236$. (For example, the case of $\sigma_{\mathrm{RS}} = 0.471$ has one of only three possible supplies randomly sampled on each growth cycle whereas $\sigma_{\mathrm{RS}} = 0.236$ has a uniform-randomly sampled resource supply. Which should truly be considered the most random or fluctuating resource supply is ambiguous.) Appendix Figure S4A shows the additional data on number of survivors as a function of number of resources that was excluded from Main Text Figure 4C. Diversity remains high until $\sigma_{\mathrm{RS}} = 0.471$, at which point diversity suddenly collapses to often only a single species surviving. This is as expected given our consideration of optimal strategies (Figure 5A-B and reproduction in Appendix Figure S4B). With only a single resource supply on each growth cycle, the optimal strategy is to be a generalist equally invested in all resources with preferences no longer mattering – i.e. the six optimal strategies suddenly "collapse" to a single strategy.

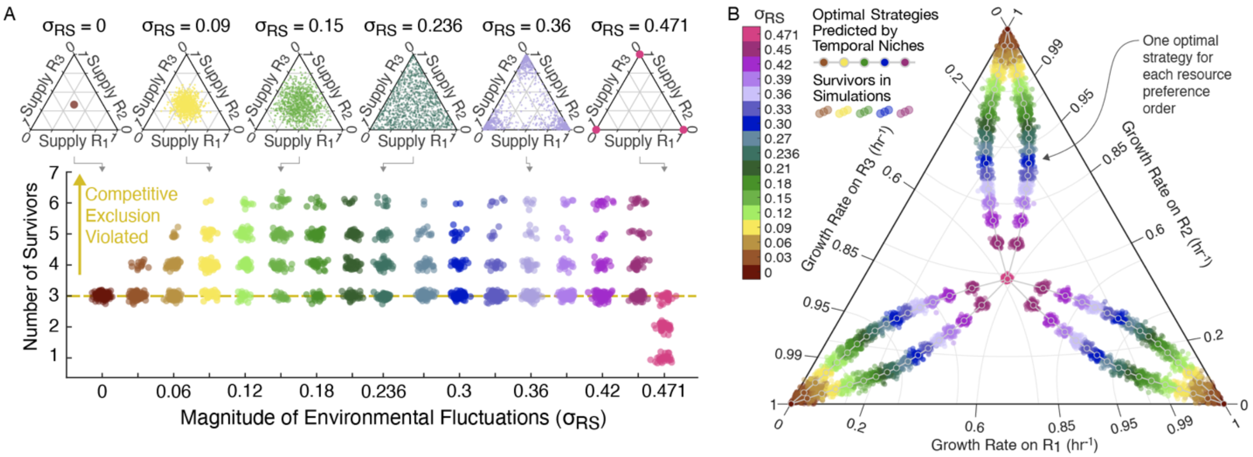


**Appendix Figure S4. Number of surviving species at all fluctuation magnitudes tested.** (**A**) Equivalent of Main Text Figure 4B-C, but now included all tested fluctuation magnitudes from $\sigma_{\mathrm{RS}} = 0$ to $\sigma_{\mathrm{RS}} = 0.471$. (**B**) Reproduction of Main Text Figure 5B to facilitate comparison to **A**.

### §2 Properties of the Diauxie Model

#### §2.1 Detailed definition of the model and importance of resource depletion times

The diauxie model we use in our paper is the same as used in the relevant Appendix sections of Bloxham et al. "Diauxic lags explain unexpected coexistence in multi-resource environments" Molecular Systems Biology 2022 and, up to a few minor changes to variable names, the model used in Wang et al. "Complementary resource preferences spontaneously emerge in diauxic microbial communities" Nature Communications 2021.

Species have order resource preferences (such as $\mathrm{R}_1 \rightarrow \mathrm{R}_3 \rightarrow \mathrm{R}_4 \rightarrow \mathrm{R}_2$) and exponential growth rates $\boldsymbol{g}_\mu = \{g_{\mu i}\}$ on each resource. Throughout, we use Greek letters ($\mu$, $\nu$, etc.) to index over species and Latin letters ($i$, $j$, etc.) to index over resources with specific species being A, B, C, etc. and specific resources being $\mathrm{R}_1$, $\mathrm{R}_2$, $\mathrm{R}_3$, etc. Species consume resources one at a time, moving to their next remaining preference when their previous is depleted. Their populations sizes are $\boldsymbol{n}(t) = \{n_\mu(t)\}$ and have dimensionless units. Variable yields and diauxic lags are not included (i.e. yields are implicitly equal to 1 and all lag times equal to 0).

The environment consists of growth and death phases (i.e. "boom-and-bust" cycles) with the combination of one growth phase and one death phase being referred to as a "growth cycle". Resources have concentrations $\boldsymbol{c}(t) = \{c_i(t)\}$. At the start of each growth cycle, resource concentrations are set to the supply concentrations $\boldsymbol{s} = \{s_i\}$. When the supply varies, we can denoted the growth cycle as a superscript to distinguish it from the resource indexing. (For example, $s_2^3$ is the supply fraction of $\mathrm{R}_2$ on growth cycle 3.) Time is reset to $t = 0$ at the start of each growth cycle. Times, population sizes, and resource concentrations can also use superscript indexing to indicate a specific growth cycle whenever dynamics over multiple cycles are being discussed. For simplicity, we assume the supply on each growth cycle is always equal to 1 (i.e. $\forall_j \sum_{i=1}^{\mathrm{N_{Re}}} s_i^j = 1$) where resource concentrations are dimensionless.

Together, the dynamic equations during the growth phase are

$$\begin{cases} \dfrac{dn_\mu}{dt} = g_{\mu\, r_\mu(\boldsymbol{c}(t))} n_\mu(t) \\ \dfrac{dc_i}{dt} = -\sum_{\mu:\, r_\mu(\boldsymbol{c}(t))=i} g_{\mu i} n_\mu(t) \end{cases},$$

where $r_\mu(\boldsymbol{c}(t))$ is equal to the index of the resource species μ is eating given resource concentrations $\boldsymbol{c}(t)$ and the notation $\mu :\, r_\mu(\boldsymbol{c}(t)) = i$ indicates a sum over all species $\mu$ that are currently consuming $\mathrm{R}_i$.

The death phase consists of dividing all population sizes by the dilution factor DF (which was always set to 10 in the Main Text). The death phase occurs as soon as $\boldsymbol{c}(t) = 0$.

As previously established (Bloxham et al., 2022), the discrete resource preferences create discrete exponential growth phases such that each species log-growth on each growth cycle is a linear function of growth rates times the resource depletion times (i.e. the times $t_{\text{dep }i}$ at which the concentration of each resource reaches zero). For example,

$$\Delta \log n_{\text{A}} = g_{\text{A1}} t_{\text{dep1}} + g_{\text{A2}}\left(t_{\text{dep2}} - t_{\text{dep1}}\right) - \log(\text{DF}) \text{ and}$$
$$\Delta \log n_{\text{B}} = g_{\text{B2}} t_{\text{dep2}} - \log(\text{DF})$$

would be the changes in population size for a species A that prefers $R_1$ over $R_2$ and a species B that prefers $R_2$ over $R_1$ on a growth cycle in which $R_1$ is depleted before $R_2$.

These equations can be rearranged to

$$\Delta \log n_{\text{A}} = (g_{\text{A1}} - g_{\text{A2}}) t_{\text{dep1}} + g_{\text{A2}} t_{\text{dep2}} - \log(\text{DF}) \text{ and}$$
$$\Delta \log n_{\text{B}} = (0) t_{\text{dep1}} + g_{\text{B2}} t_{\text{dep2}} - \log(\text{DF}) ,$$

or in matrix form

$$\begin{pmatrix} \Delta \log n_{\text{A}} \\ \Delta \log n_{\text{B}} \end{pmatrix} = \begin{pmatrix} g_{\text{A1}} - g_{\text{A2}} & g_{\text{A2}} \\ 0 & g_{\text{B2}} \end{pmatrix} \begin{pmatrix} t_{\text{dep1}} \\ t_{\text{dep2}} \end{pmatrix} - \log(\text{DF}),$$

and in general can be written in the form

$$\Delta \log \boldsymbol{n} = \mathbf{G}(\text{DO})\, \boldsymbol{t}_{\text{dep}} - \log(\text{DF}) ,$$

where $\mathbf{G}(\text{DO})$ depends on the "depletion order" DO (i.e. the order in which resources are depleted) and has terms of the forms $g_{\mu i}$, $(g_{\mu i} - g_{\mu j})$, and 0.

Species' growth and resource consumption continuously affect $\boldsymbol{c}(t)$, which determine $\boldsymbol{t}_{\text{dep}}$ which in turn determines $\mathbf{G}(\text{DO})$, but the times $\boldsymbol{t}_{\text{dep}}$ fully determine each species' log-growth on each growth cycle. Thus, the resource depletion times $\boldsymbol{t}_{\text{dep}}$ mediate all interspecies interactions, making them central to all dynamics.

##### §2.2 At a fixed point, competitive exclusion is obeyed

We define a fixed point as having population sizes that do not vary between growth cycles, and assume a constant resource supply whenever we're discussing fixed points. The system has the same initial condition $\{\boldsymbol{n}, \boldsymbol{c}\}$ at the start of each growth cycle therefore the same $\boldsymbol{t}_{\text{dep}}$ on each growth cycle. Constant population sizes require

$$0 = \mathbf{G}(\text{DO})\, \boldsymbol{t}_{\text{dep}} - \log(\mathbf{DF}) .$$

As previously established (Bloxham et al. 2022) and excluding the case of fine-tuned growth rates, this equation can only be solved if there are fewer or an equal number of surviving species as resources (i.e. if $N_{\text{Sp}} \leq N_{\text{Re}}$ where $N_{\text{Sp}}$ is the number of surviving species and $N_{\text{Re}}$ is the number of resources). This is the "competitive exclusion principle".

Species population sizes affect $\boldsymbol{t}_{\text{dep}}$ which then determines the change in population sizes on each growth cycle. Depending on species' growth rates and resource preferences, this feedback can bring the system to a stable fixed point with $N_{\text{Sp}} \leq N_{\text{Re}}$.

Population sizes affect the depletion order and therefore which $\mathbf{G}(\text{DO})$ through its DO dependence but each $\mathbf{G}(\text{DO})$ can only have a discrete set of values (one for each resource depletion order) and there are still only $N_{\text{Re}}$ degrees of freedom, so this does not create the necessary flexibility to break the competitive exclusion principle.

##### §2.3 With fluctuations, exponentially many species as resources can coexist

By "fluctuations" we mean any dynamics in which population sizes do not reach a fixed point but instead vary from one growth cycle to the next. This includes periodic oscillations and chaos under a fixed resource supply and the case of environmental fluctuations (e.g. a resource supply $\boldsymbol{s}$ that varies between growth cycles).

When average over all growth cycles, all surviving species must obey

$$\langle \Delta \log n_\mu \rangle = 0 \,,$$

or else the species' population size would be going to infinity (which is impossible under a finite resource supply and dilution factor) or to zero (in which case we would consider the species extinct).

###### *§2.3.1 An initial approach via a summation over depletion orders*

If the depletion order were constant, we would have

$$0 = \mathbf{G}(\mathrm{DO})\, \langle \boldsymbol{t}_{\mathrm{dep}} \rangle - \log(\mathbf{DF}) \,,$$

in which case there would still only be $N_{\mathrm{Re}}$ free variables $\{\langle \boldsymbol{t}_{\mathrm{dep}\,i} \rangle\}$ and competitive exclusion would still be obeyed.

However, fluctuating population sizes can lead to a variable depletion order, which would expand the zero-average-growth equation to

$$0 = \sum_{i=1}^{N_{\mathrm{Re}}!} \mathbf{G}(\mathrm{DO}_i)\, \langle \boldsymbol{t}_{\mathrm{dep}} \rangle_{\mathrm{DO}_i} - \log(\mathbf{DF})$$

where $\langle \boldsymbol{t}_{\mathrm{dep}} \rangle_{\mathrm{DO}\,i}$ indicates the average $\boldsymbol{t}_{\mathrm{dep}}$ when the depletion order is $\mathrm{DO}_i$ (with $N_{\mathrm{Re}}!$ possible depletion orders) multiplied by the frequency of that depletion order occurring. To turn this into a matrix equation, the $\mathbf{G}(\mathrm{DO}_i)$ could be concatenated horizontally and the $\langle \boldsymbol{t}_{\mathrm{dep}} \rangle_{\mathrm{DO}_i}$ vertically. This yields a massive combined $\langle \boldsymbol{t}_{\mathrm{dep}} \rangle_{\mathrm{DO}_i}$ vector of length $N_{\mathrm{Re}} * N_{\mathrm{Re}}!$, implying that $N_{\mathrm{Re}} * N_{\mathrm{Re}}!$ species might coexist. However, the combined matrix of stacked $\mathbf{G}(\mathrm{DO}_i)$ will have enough repeating values and patterns to never reach rank-($N_{\mathrm{Re}} * N_{\mathrm{Re}}!$) (or even close to it for moderate to large $N_{\mathrm{Re}}$). Indeed, a much tighter bound exists if we take a distinctly different and novel approach to understanding this system, and this alternative approach will grant us much better insights into the dynamics of this system when population sizes fluctuate.

###### *§2.3.2 A tighter exponential bound and better understanding via temporal niches*

For a better approach to this system, we define the concept of a "temporal niche", which is simply a specific combination of which resources have not yet been depleted. For example, all resources being present is a temporal niche, as is only $R_1$ being present or both $R_2$ and $R_3$ being present.

For further illustration, when the depletion order is $t_{\mathrm{dep1}} < t_{\mathrm{dep2}} < t_{\mathrm{dep3}}$ species experience the $R_1 + R_2 + R_3$ niche then the $R_2 + R_3$ niche then the $R_3$ niche. When the depletion order is $t_{\mathrm{dep2}} < t_{\mathrm{dep1}} < t_{\mathrm{dep3}}$ species experience the $R_1 + R_2 + R_3$ niche then the $R_1 + R_3$ niche then the $R_3$ niche. Under each of these depletion orders, the first niche experienced is $R_1 + R_2 + R_3$ and the last niche experienced is $R_3$. So, whenever the depletion order is $t_{\mathrm{dep1}} < t_{\mathrm{dep2}} < t_{\mathrm{dep3}}$ or $t_{\mathrm{dep2}} < t_{\mathrm{dep1}} < t_{\mathrm{dep3}}$, species end the growth phase growing on $R_3$ and at their $g_{\mu 3}$ growth rates. Importantly, growth rates are the same every time a temporal niche occurs regardless of what order resources were depleted in to get to that temporal niche.

Understanding this system from the temporal niche perspective is a distinct approach from the $\mathbf{G}(\mathrm{DO})$ approach explored above and will involve constructing an alternative matrix $\widetilde{\mathbf{G}}$ that contains species' growth rates arranged by temporal niche.

The zero-average-growth equation can now be expressed as

$$0 = \widetilde{\mathbf{G}}\,\langle \boldsymbol{t}_{\mathrm{tn}} \rangle - \log(\mathbf{DF})\,,$$

where $\widetilde{\mathbf{G}}$ contains species' growth rates by temporal niche and $\langle \boldsymbol{t}_{\mathrm{tn}} \rangle$ is the average time spent in each temporal niche. For example, consider three species competing for two resources with species A and B preferring $R_1$ over $R_2$ and species C preferring $R_2$ over $R_1$. In the $R_1 + R_2$ niche, species A and B are eating $R_1$ and growing at their $R_1$ growth rates while species C is eating $R_2$ and growing at its $R_2$ growth rate. In the $R_1$ only and $R_2$ only niches all species are eating $R_1$ and $R_2$, respectively. Therefore, for this example

$$\widetilde{\mathbf{G}} = \begin{pmatrix} g_{\mathrm{A1}} & g_{\mathrm{A1}} & g_{\mathrm{A2}} \\ g_{\mathrm{B1}} & g_{\mathrm{B1}} & g_{\mathrm{B2}} \\ g_{\mathrm{C2}} & g_{\mathrm{C1}} & g_{\mathrm{C2}} \end{pmatrix},$$

and the zero-average-growth equation is

$$0 = \begin{pmatrix} g_{\mathrm{A1}} & g_{\mathrm{A1}} & g_{\mathrm{A2}} \\ g_{\mathrm{B1}} & g_{\mathrm{B1}} & g_{\mathrm{B2}} \\ g_{\mathrm{C2}} & g_{\mathrm{C1}} & g_{\mathrm{C2}} \end{pmatrix} \begin{pmatrix} \langle t_{12} \rangle \\ \langle t_1 \rangle \\ \langle t_2 \rangle \end{pmatrix} - \log(\mathbf{DF})\,.$$

For six species competing for three resources with resource preferences as defined in the below table the zero-average-growth equation containing the matrix $\widetilde{\mathbf{G}}$ would be:

| | 1st | 2nd | 3rd |
|---|---|---|---|
| A | $R_1$ | $R_2$ | $R_3$ |
| B | $R_1$ | $R_3$ | $R_2$ |
| C | $R_2$ | $R_1$ | $R_3$ |
| D | $R_2$ | $R_3$ | $R_1$ |
| E | $R_3$ | $R_1$ | $R_2$ |
| F | $R_3$ | $R_2$ | $R_1$ |

$$0 = \begin{pmatrix} g_{\mathrm{A1}} & g_{\mathrm{A1}} & g_{\mathrm{A1}} & g_{\mathrm{A2}} & g_{\mathrm{A1}} & g_{\mathrm{A2}} & g_{\mathrm{A3}} \\ g_{\mathrm{B1}} & g_{\mathrm{B1}} & g_{\mathrm{B1}} & g_{\mathrm{B3}} & g_{\mathrm{B1}} & g_{\mathrm{B2}} & g_{\mathrm{B3}} \\ g_{\mathrm{C2}} & g_{\mathrm{C2}} & g_{\mathrm{C1}} & g_{\mathrm{C2}} & g_{\mathrm{C1}} & g_{\mathrm{C2}} & g_{\mathrm{C3}} \\ g_{\mathrm{D2}} & g_{\mathrm{D2}} & g_{\mathrm{D3}} & g_{\mathrm{D2}} & g_{\mathrm{D1}} & g_{\mathrm{D2}} & g_{\mathrm{D3}} \\ g_{\mathrm{E3}} & g_{\mathrm{E1}} & g_{\mathrm{E3}} & g_{\mathrm{E3}} & g_{\mathrm{E1}} & g_{\mathrm{E2}} & g_{\mathrm{E3}} \\ g_{\mathrm{F3}} & g_{\mathrm{F2}} & g_{\mathrm{F3}} & g_{\mathrm{F3}} & g_{\mathrm{F1}} & g_{\mathrm{F2}} & g_{\mathrm{F3}} \end{pmatrix} \begin{pmatrix} \langle t_{123} \rangle \\ \langle t_{12} \rangle \\ \langle t_{13} \rangle \\ \langle t_{23} \rangle \\ \langle t_1 \rangle \\ \langle t_2 \rangle \\ \langle t_3 \rangle \end{pmatrix} - \log(\mathbf{DF})$$

The 42 entries in this 6x7 example $\widetilde{\mathbf{G}}$ matrix only have 18 unique values, but because they are arranged differently in each column, this remains a rank-6 matrix (and would be a rank-7 matrix if a seventh species were added).

There are $2^{N_{\text{Re}}}-1$ temporal niches (all the binary combinations of whether or not a resource is present minus 1 for the case of no resources remaining), so, to our current understanding, $N_{\text{Sp}} \leq 2^{N_{\text{Re}}}-1$ becomes the new upper bound on the number surviving species.

Each resource preference order distributes the species' growth rates differently across the temporal niches, and if each species has a different resource preference order the matrix $\widetilde{\mathbf{G}}$ will have no rank-degeneracies. It is also possible for coexisting species to share a resource preference order, including when diversity is saturated at $N_{\text{Sp}} = 2^{N_{\text{Re}}}-1$ (as is the case in the Appendix *§3.5.1* and *§3.5.2* examples). However, it is not possible for more surviving species than resources to share the same depletion order or else $\widetilde{\mathbf{G}}$ becomes degenerate and it is no longer possible to find a $\langle \boldsymbol{t}_{\text{tn}} \rangle$ that lets all those species survive.

#### §2.4 Optimal strategies

If $\langle \boldsymbol{t}_{\text{tn}} \rangle$ is known then $\langle \Delta \log n_\mu \rangle = \widetilde{\mathbf{G}}_\mu \langle \boldsymbol{t}_{\text{tn}} \rangle - \log(\mathbf{DF})$ can be calculated for all competing species. With a constraint such as $\sum_{i=1}^{N_{\text{Re}}} {g_{\mu i}}^2 = 1\,\text{hr}^{-2}$, a species growth rates can be optimized to maximize $\langle \Delta \log n_\mu \rangle$. To determine optimal strategies for comparison to the growth rates of the survivors, we took $\langle \boldsymbol{t}_{\text{tn}} \rangle$ from the simulation data (averaging across all simulations for fluctuation magnitude) and calculated the optimal $\boldsymbol{g}_\mu$ for each resource preference order (the quadratic form of the constraint making this optimization nothing more than the renormalization of a vector). Symmetry of the resource supply meant that each optimal strategy had the same first, second, and third preference growth rates, which were then distributed into $\boldsymbol{g}$ according to the species' resource preference orders.

### §3 Additional Results with a Constant Resource Supply

#### §3.1 With three species competing for two resources a fixed point was always reached

We randomly sampled 4*$10^9$ communities of three species competing for two resources. We ensured that in each random community both resource preference orders were represented. Growth rates were log-uniform randomly sampled from $10^{-4}$ to 1 $hr^{-1}$. The dilution factor (equal to 10 in the Main Text) was log-uniform randomly sampled from 2 to 25, and the (constant) resource supply fractions were uniform randomly sampled from $\{s_1, s_2\} = \{0,1\}$ to $\{s_1, s_2\} = \{1,0\}$. In none of the communities did an oscillation occur. Given the massive number of random communities sampled, we believe a three-species oscillation with only two resources supplied in constant fractions is not possible. Two species cannot oscillate, and we have already shown that four or more species cannot survive on only two resources. Therefore, oscillations on only two resources supplied in constant amounts are not possible.

#### §3.2 When sampling random communities, emergent oscillations are rare

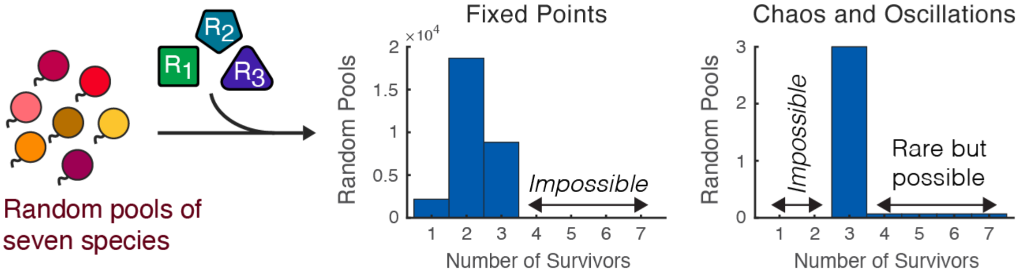


**Appendix Figure S5. Emergent oscillations are rare when sampling random species.** We sampled random pools of 7 species competing for 3 resources. The dilution factor was 10 and the resource supply was constant with equal fractions of each resource. Growth rates were sampled from the same $\sum_{i=1}^{N_{Re}} g_{\mu i}^{2} = 1\ \mathrm{hr}^{-2}$ distribution as in the simulations with a fluctuating resource supply. We sorted by whether the community reached a steady-state oscillation or chaos and by number of survivors. Each community was started at 36 different initial species fractions and was categorized as oscillating or being chaotic if any of the initial conditions resulted in that. For number of survivors, the greatest number from each of the initial conditions was used. After testing approximately $3 * 10^4$ random communities, we observed 3 communities that oscillated.

#### §3.3 Example of five species coexisting on four resources without anomalous resource preferences

"Anomalous resource preferences" refers to a species preferring a resource it grows slower on over a resource it grows faster on (i.e. resource preferences not matching the ordering of the species' growth rates). The example presented in Main Text Figure 1C and explored in Main Text Figure 2 had anomalous resource preferences, potentially raising the question of whether anomalous resource preference is necessary for oscillations and competitive exclusion violations in the case of a constant resource supply. Appendix Figure S6 presents an example demonstrating that this is not the case.

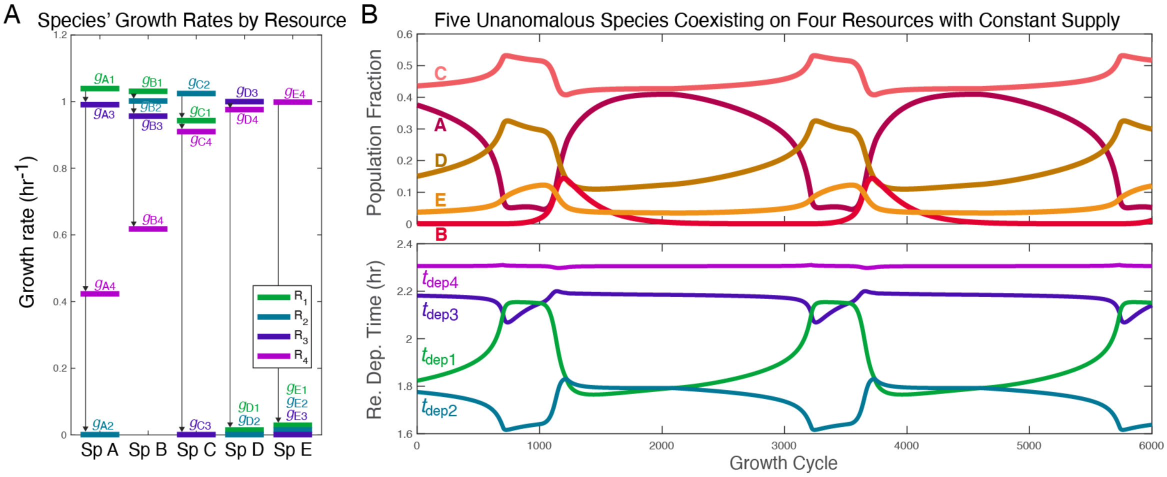


**Appendix Figure S6. Example of an oscillation and competitive exclusion violation without anomalous resource preferences.** (**A**) Growth rates for the five species on each of the four resources in this example. Growth rates are indicated by horizontal bars, colored by resource and also labeled. Arrows illustrate the species' resource preference orders, which always match their descending growth rate orders. (**B**) The steady-state oscillation in which the four species coexist, showing population fractions on top and resource depletion times on the bottom.

In this example, the dilution factor is 10, the resource supply fractions are

$$\boldsymbol{s} = \{0.25, 0.25, 0.25, 0.25\},$$

and species growth rates and resource preferences are given by Appendix Table S12.

| | Growth Rates (hr$^{-1}$) | | | | Preferences | | | |
|---|---|---|---|---|---|---|---|---|
| | **$R_1$** | **$R_2$** | **$R_3$** | **$R_4$** | **1$^{st}$** | **2$^{nd}$** | **3$^{rd}$** | **4$^{th}$** |
| **Sp. A** | 1.0392 | 0.0010 | 0.9908 | 0.4232 | R1 | R3 | R4 | R2 |
| **Sp. B** | 1.0312 | 1.0020 | 0.9565 | 0.6177 | R1 | R2 | R3 | R4 |
| **Sp. C** | 0.9432 | 1.0241 | 0.0010 | 0.9102 | R2 | R1 | R4 | R3 |
| **Sp. D** | 0.0150 | 0.0010 | 1.0001 | 0.9759 | R3 | R4 | R1 | R2 |
| **Sp. E** | 0.0290 | 0.0150 | 0.0010 | 0.9987 | R4 | R1 | R2 | R3 |

**Appendix Table S12.** Growth rates and resource preferences for the example of an oscillation and competitive exclusion violation without anomalous resource preferences (Appendix Figure S6).

#### §3.4 Example of chaos

Only periodic oscillations were presented in the Main Text. Diauxic growth does, however, exhibit chaos, as is demonstrated in Appendix Figure S7.

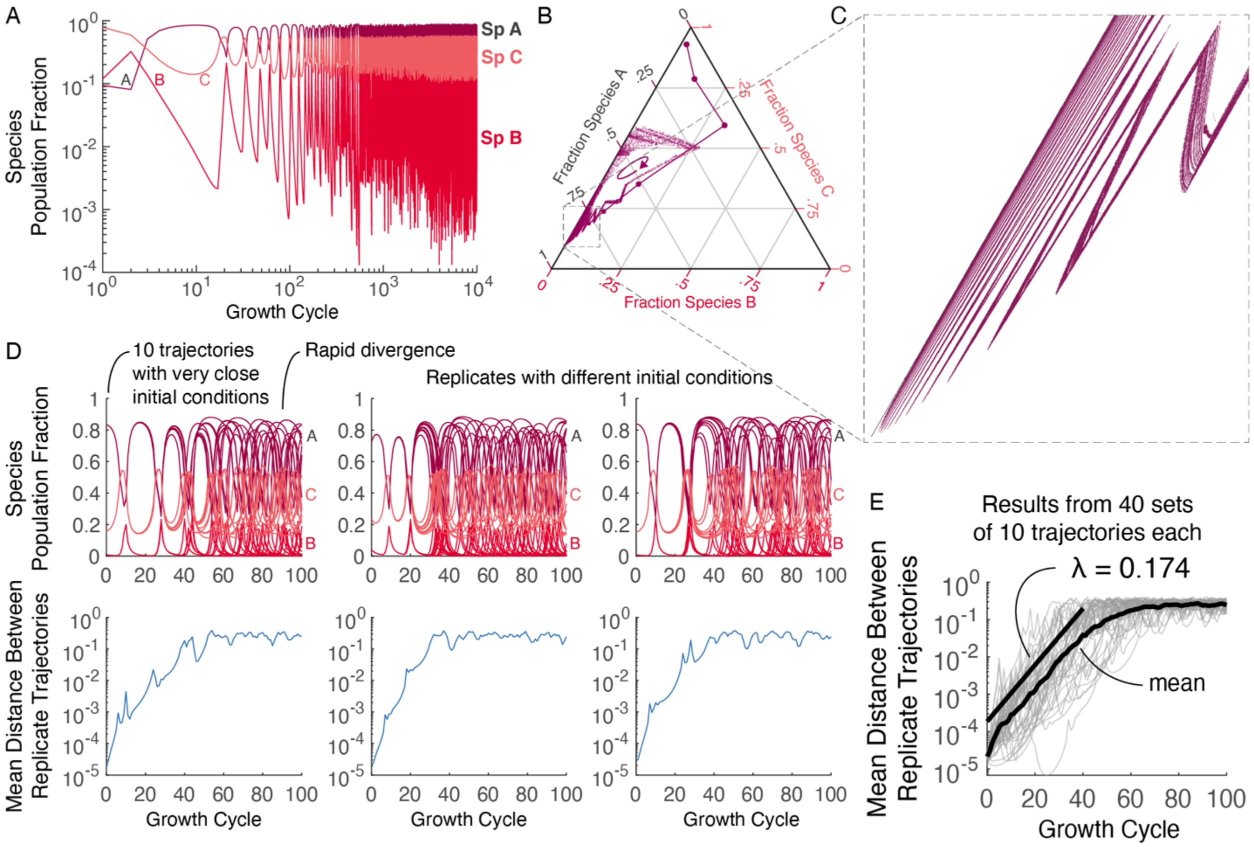


**Appendix Figure S7. Diauxic growth can exhibit chaos.** (**A**) Population fractions at the end of each of $10^4$ growth cycles for a three-species community with a chaotic attractor. Simulation parameters are presented in Appendix Table S12. (**B**) The same trajectory as in **A** plotted on a simplex diagram, with population fraction fractions mapped onto the simplex diagram in the same manner resource supply fractions were mapped onto their simplex diagrams (Appendix §1.2 and Figure S1). The inoculum condition (highest point on the simplex) and compositions after the first five growth cycles are shown as enlarged, connected dots. Community compositions for the remaining 9,995 growth cycles are shown as small, unconnected dots. Curved arrow illustrates direction of motion within the chaotic attractor. It takes $16.7 \pm 4.4$ growth cycles (mean and standard deviation) to travel around the attractor once. (**C**) Magnification of the region indicated by the box overlay on **B**. The simulation was run for $2 * 10^6$ growth cycles, the last $10^6$ of which are shown here. A fractal-like pattern – as is often associated with chaos – is evident. (**D**) To rigorously established chaos, we randomly chose 40 points from the trajectory and around each of these points created a cluster of 10 initial conditions by adding normal random noise with $\sigma = 10^{-5}$ to each species' population fraction and then renormalizing the population fractions to $\sum_{\mu=1}^{3} f_\mu = 1$. We simulated the community starting from each of the 40 sets of 10 initial conditions for 100 growth cycles. Each column of **D** shows one of the 40 sets of initial conditions. In each column, the top plot shows the trajectories for the 10 closely clustered initial conditions and the bottom plot shows the mean L2-distance between trajectories. Trajectories can be seen to rapidly diverge in the top plot as is confirmed and quantified in the bottom plots. (**E**) Light grey lines show the mean L2-distances between trajectories for each of the 40 clusters of initial conditions. The black curve is the mean of these means (in log-space). The straight black line depicts the slope extracted from this mean over the first 40 growth cycles. It has a slope of $\lambda = 0.174 \pm 0.002$ (on a natural log scale). This slope represents an approximate Lyapunov exponent for the chaotic attractor.

In the example of a chaotic attractor presented in Appendix Figure S7, the dilution factor is 18.5, the resource supply is

$$\boldsymbol{s} = \{0.35, 0.01, 0.55\}\,,$$

and the species' growth rates and resource preferences are given by Appendix Table S13.

| | **Growth Rates ($hr^{-1}$)** | | | **Preferences** | | |
|---|---|---|---|---|---|---|
| | **$R_1$** | **$R_2$** | **$R_3$** | **1st** | **2nd** | **3rd** |
| **Sp. A** | 0.5100 | 0.0400 | 0.8400 | R2 | R3 | R1 |
| **Sp. B** | 0.0350 | 0.0950 | 0.7550 | R2 | R1 | R3 |
| **Sp. C** | 0.1000 | 0.0335 | 0.1800 | R1 | R2 | R3 |

**Appendix Table S13.** Growth rates and resource preferences for the example of chaos in Appendix Figure S7.

#### §3.5 Examples of seven species on three resources, fifteen on four, and twenty-three on five

##### *§3.5.1 Seven Species on Three Resources*

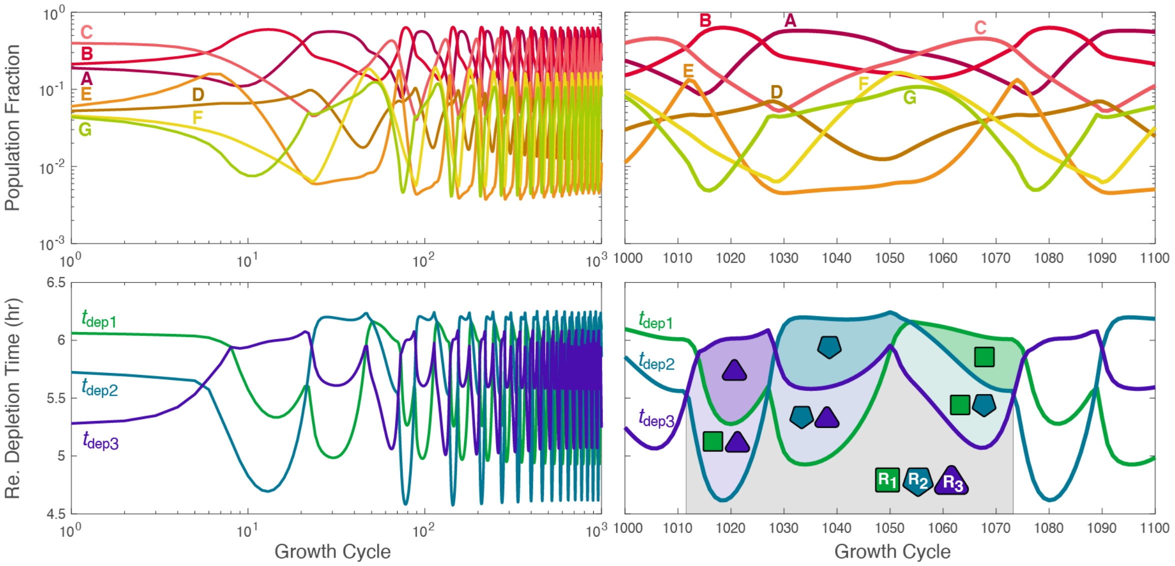


**Appendix Figure S8. Seven species stably coexisting on three resources.** Species population fractions (top) and resource depletion times (bottom) over the first 1000 growth cycles (left) and growth cycles 1000 through 1100 (right). In the bottom right plot, the seven temporal niches are highlighted as in Main Text Figure 2C. All seven species coexist, saturating the upper bound. In this simulation, the dilution factor is 10, the resource supply is constant with an equal supply of all three resources, and species' growth rates and preferences are given by Appendix Table S14.

| | Growth Rates ($hr^{-1}$) | | | Preferences | | |
|---|---|---|---|---|---|---|
| | **$R_1$** | **$R_2$** | **$R_3$** | **1st** | **2nd** | **3rd** |
| **Sp. A** | 0.3705 | 0.0209 | 0.7063 | R1 | R3 | R2 |
| **Sp. B** | 0.7602 | 0.3679 | 0.1326 | R2 | R1 | R3 |
| **Sp. C** | 0.3222 | 0.5735 | 0.3658 | R3 | R2 | R1 |
| **Sp. D** | 0.3860 | 0.0376 | 0.3881 | R3 | R1 | R2 |
| **Sp. E** | 1.0273 | 0.3740 | 0.2214 | R2 | R3 | R1 |
| **Sp. F** | 0.3364 | 0.7009 | 0.3690 | R3 | R1 | R2 |
| **Sp. G** | 0.0165 | 0.3777 | 0.9944 | R2 | R1 | R3 |

**Appendix Table S14.** Growth rates and resource preferences for the example of seven species coexisting on three resources in Appendix Figure S8.

*§3.5.2 Fifteen Species on Four Resources*

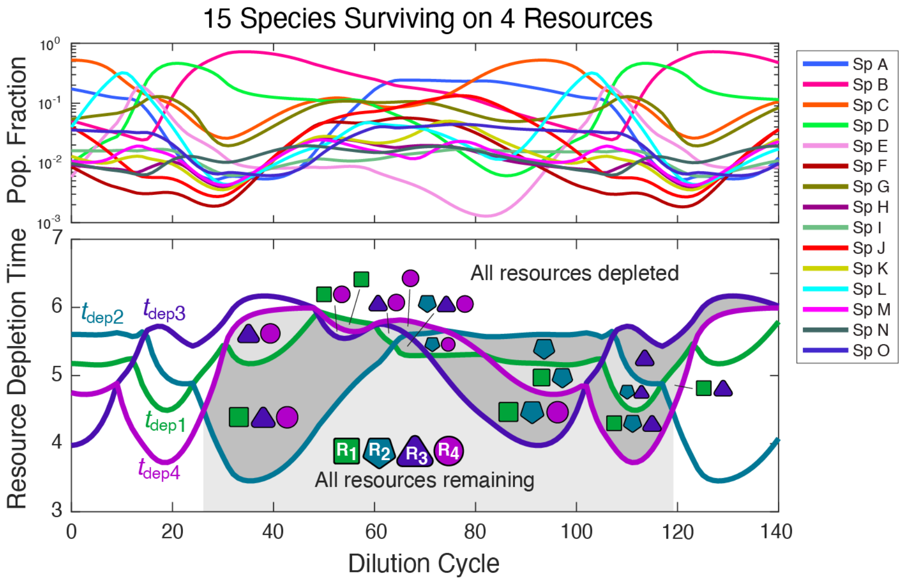


**Appendix Figure S9.** Species population fractions (top) and resource depletion times (bottom) over the 140 growth cycles after the community has settled into its steady-state oscillation. In the bottom plot, the fifteen temporal niches are highlighted as in Main Text Figure 2C. (The temporal niche in which $R_3$ and $R_4$ are the two remaining resource occurs twice.) All fifteen species coexist, saturating the upper bound. In this simulation, the dilution factor is 10, the resource supply is constant with an equal supply of all three resources, and species' growth rates and preferences are given by Appendix Table S15.

| | Growth Rates ($hr^{-1}$) | | | | Preferences | | | |
|---|---|---|---|---|---|---|---|---|
| | **$R_1$** | **$R_2$** | **$R_3$** | **$R_4$** | **1st** | **2nd** | **3rd** | **4th** |
| **Sp. A** | 0.4297 | 0.1003 | 0.0644 | 0.0543 | R1 | R4 | R3 | R2 |
| **Sp. B** | 0.3210 | 0.4014 | 0.9676 | 0.1506 | R2 | R1 | R4 | R3 |
| **Sp. C** | 0.2344 | 0.3529 | 0.3911 | 0.6520 | R3 | R4 | R1 | R2 |
| **Sp. D** | 0.8959 | 0.5339 | 0.0080 | 0.3826 | R4 | R1 | R2 | R3 |
| **Sp. E** | 0.7888 | 0.8726 | 0.0172 | 0.3854 | R4 | R1 | R3 | R2 |
| **Sp. F** | 0.1405 | 0.3597 | 0.4057 | 0.1178 | R3 | R2 | R1 | R4 |
| **Sp. G** | 0.3257 | 0.5155 | 0.2032 | 0.3978 | R4 | R2 | R3 | R1 |
| **Sp. H** | 0.3921 | 0.5906 | 0.4136 | 0.3241 | R1 | R2 | R3 | R4 |
| **Sp. I** | 0.3432 | 0.4110 | 0.3418 | 0.0983 | R2 | R3 | R4 | R1 |
| **Sp. J** | 0.3954 | 0.1082 | 0.4022 | 0.4889 | R3 | R4 | R1 | R2 |
| **Sp. K** | 0.2547 | 0.8264 | 0.3963 | 0.7528 | R3 | R1 | R4 | R2 |
| **Sp. L** | 1.2296 | 0.3677 | 0.2989 | 0.3937 | R4 | R3 | R1 | R2 |
| **Sp. M** | 0.2403 | 0.9877 | 0.3938 | 0.1555 | R2 | R1 | R2 | R4 |
| **Sp. N** | 0.3034 | 0.4053 | 0.5830 | 0.3321 | R2 | R4 | R1 | R3 |
| **Sp. O** | 0.4258 | 0.4096 | 0.2134 | 0.1087 | R2 | R1 | R4 | R3 |

**Appendix Table S15.** Growth rates and resource preferences for the example of fifteen species coexisting on four resources in Appendix Figure S9.

##### *§3.5.3 Twenty-Three Species on Five Resources*

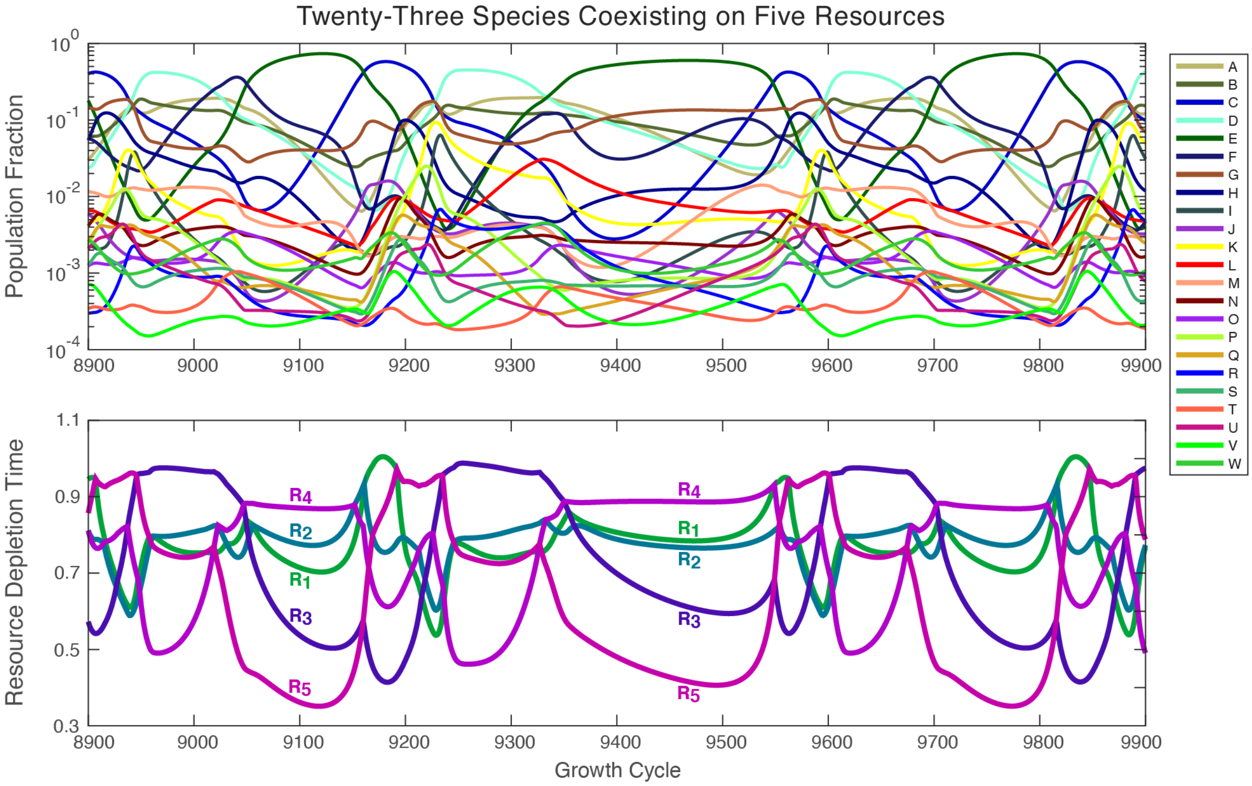


**Appendix Figure S10.** Species population fractions (top) and resource depletion times (bottom) over 1000 growth cycles after the community has settled into its steady-state oscillation.

In this example, the dilution factor is 1.8167, the resource supply fractions are

$$\boldsymbol{s} = \{0.1869, 0.1945, 0.2067, 0.1974, 0.2145\}\,,$$

and species growth rates and resource preferences are given by Appendix Table S16. All twenty three species coexist.

| | Growth Rates ($hr^{-1}$) | | | | | Preferences | | | | |
|---|---|---|---|---|---|---|---|---|---|---|
| | $R_1$ | $R_2$ | $R_3$ | $R_4$ | $R_5$ | 1st | 2nd | 3rd | 4th | 5th |
| **Sp. A** | 0.6731 | 0.1190 | 0.4319 | 0.5263 | 0.8382 | R1 | R3 | R4 | R2 | R5 |
| **Sp. B** | 0.3896 | 0.6748 | 0.3016 | 0.6088 | 0.6590 | R2 | R4 | R1 | R5 | R3 |
| **Sp. C** | 0.3606 | 0.5542 | 0.5965 | 0.9641 | 0.2867 | R3 | R4 | R2 | R5 | R1 |
| **Sp. D** | 0.1393 | 0.4955 | 0.2140 | 0.6611 | 0.8378 | R4 | R5 | R1 | R2 | R3 |
| **Sp. E** | 0.6158 | 0.9342 | 0.9882 | 0.6559 | 0.5534 | R5 | R3 | R1 | R2 | R4 |
| **Sp. F** | 0.6735 | 0.5056 | 0.8929 | 1.6611 | 0.6086 | R5 | R2 | R1 | R4 | R4 |
| **Sp. G** | 0.1745 | 0.7118 | 0.1292 | 0.4770 | 0.5150 | R2 | R1 | R4 | R3 | R5 |
| **Sp. H** | 0.7112 | 0.6236 | 0.1181 | 0.3825 | 0.3221 | R1 | R5 | R4 | R2 | R3 |
| **Sp. I** | 0.6111 | 0.9918 | 0.3287 | 1.3090 | 0.9581 | R1 | R2 | R4 | R3 | R5 |
| **Sp. J** | 0.6919 | 0.9312 | 0.6146 | 0.7734 | 0.5631 | R3 | R5 | R2 | R4 | R1 |
| **Sp. K** | 0.6350 | 0.9367 | 0.3415 | 0.9948 | 1.1029 | R1 | R2 | R3 | R5 | R4 |
| **Sp. L** | 0.7858 | 0.9690 | 0.5843 | 0.7131 | 0.6176 | R5 | R4 | R2 | R1 | R3 |

| | | | | | | | | | |
|---|---|---|---|---|---|---|---|---|---|---|
| **Sp. M** | 0.6762 | 0.5991 | 0.6157 | 1.3539 | 0.7454 | R3 | R2 | R1 | R5 | R4 |
| **Sp. N** | 0.6736 | 0.5820 | 0.4385 | 0.6561 | 0.3216 | R1 | R3 | R4 | R5 | R2 |
| **Sp. O** | 0.6470 | 0.3125 | 0.6237 | 0.6665 | 0.4915 | R4 | R5 | R3 | R1 | R2 |
| **Sp. P** | 0.6912 | 0.6477 | 0.2397 | 0.5312 | 0.7156 | R1 | R3 | R2 | R5 | R4 |
| **Sp. Q** | 0.6836 | 0.0148 | 0.2540 | 0.6582 | 0.3759 | R1 | R4 | R5 | R3 | R2 |
| **Sp. R** | 0.6324 | 0.9637 | 0.4105 | 0.9099 | 0.9180 | R1 | R4 | R2 | R3 | R5 |
| **Sp. S** | 0.6740 | 0.2194 | 0.4359 | 0.6625 | 0.3056 | R1 | R3 | R4 | R5 | R2 |
| **Sp. T** | 0.6037 | 0.7029 | 0.6492 | 0.6664 | 0.4719 | R4 | R5 | R1 | R2 | R3 |
| **Sp. U** | 0.6645 | 0.6366 | 0.3114 | 0.8703 | 0.5693 | R1 | R2 | R3 | R4 | R5 |
| **Sp. V** | 1.0298 | 0.3915 | 0.7212 | 0.7866 | 0.5960 | R5 | R3 | R4 | R2 | R1 |
| **Sp. W** | 0.8099 | 0.1248 | 0.6297 | 0.7861 | 0.5512 | R3 | R5 | R4 | R1 | R2 |

**Appendix Table S16.** Growth rates and resource preferences for the example of twenty three species coexisting on five resources in Appendix Figure S10.

#### §3.6 Intuitive Construction of an Oscillating Community

In this appendix section, we demonstrate the intuitive construction of oscillating communities with more coexisting species than resources and comment along the way on methodology for producing examples that maximize the number of coexisting species. These intuitive construction rely heavily on anomalous resource preference (that don't match the ordering of species' growth rate) and species being cyclic permutations of each other, but neither of these properties is necessary.

##### *§3.6.1 Constructing an Oscillation of Three Species on Three Resources*

To construct an oscillating community of three species competing for three resources, we can, for simplicity, assume the species are cyclic permutations of each other such species A prefers R1 then R2 then R3, species B prefers R2 then R3 then R1, and species C prefers R3 then R1 then R2. All species will have the same first-, second-, and third-preference growth rates (i.e. $g_{A1} = g_{B2} = g_{C3} = g_{1^{st}}$, $g_{A2} = g_{B3} = g_{C1} = g_{2^{nd}}$, and $g_{A3} = g_{B1} = g_{C2} = g_{3^{rd}}$).

For an oscillation to occur, we will want species A having a large population size to promote the growth of B's population size and the shrinking of C's population. Because species A consumes R1 then R2 then R3, when species A is most of the population resources will be depleted in this order and the all-resource niche, the R2 and R3 niche and the R3 only niche will occur. Species B will be growing on its first preference, R2, during these first two niches and on its second preference, R3, during the last niche. Meanwhile, species C will be growing on its first preference, R3, in all these niches. So to have species B's population grow while species C's population shrinks will require $g_{2^{nd}} > g_{1^{st}}$.

As we continue to think through our construction of this oscillation, we also note that we will want species A's population size to begin to shrink as the size of B's population starts to increase significantly. Species A consumes R1 first and B consumes R2 first and both at the same rate, so when the population is dominated by roughly equal populations of A and B, R1 and R2 will be depleted at roughly the same time and the all-resource niche and the R3-only niche will dominate the dynamics. To have species A's population size be decreasing at this point will require it to have a slow R3 growth rate. This leads us to the growth rate ordering $g_{2^{nd}} > g_{1^{st}} > g_{3^{rd}}$.

In Appendix Figure S11, we explore the oscillation that is created between species with resource preferences and growth rates

| | **Growth Rates (hr$^{-1}$)** | | | **Preferences** | | |
|---|---|---|---|---|---|---|
| | **$R_1$** | **$R_2$** | **$R_3$** | **1$^{st}$** | **2$^{nd}$** | **3$^{rd}$** |
| **Sp. A** | 0.6–δg | 0.6+δg | 0.2 | R1 | R2 | R3 |
| **Sp. B** | 0.2 | 0.6–δg | 0.6+δg | R3 | R1 | R2 |
| **Sp. C** | 0.6+δg | 0.2 | 0.6–δg | R2 | R3 | R1 |

(consistent with the above discussion) as δg is increased. As expected, as δg becomes sufficiently large the community experiences a Hopf bifurcation and begins to oscillate. A small amount of random noise can be added to the growth rates without losing the oscillation (Appendix Figure S13), confirming true rather than neutral stability.

This oscillation is essentially the same setup as the one briefly discussed in the Appendix to Wang 2021.

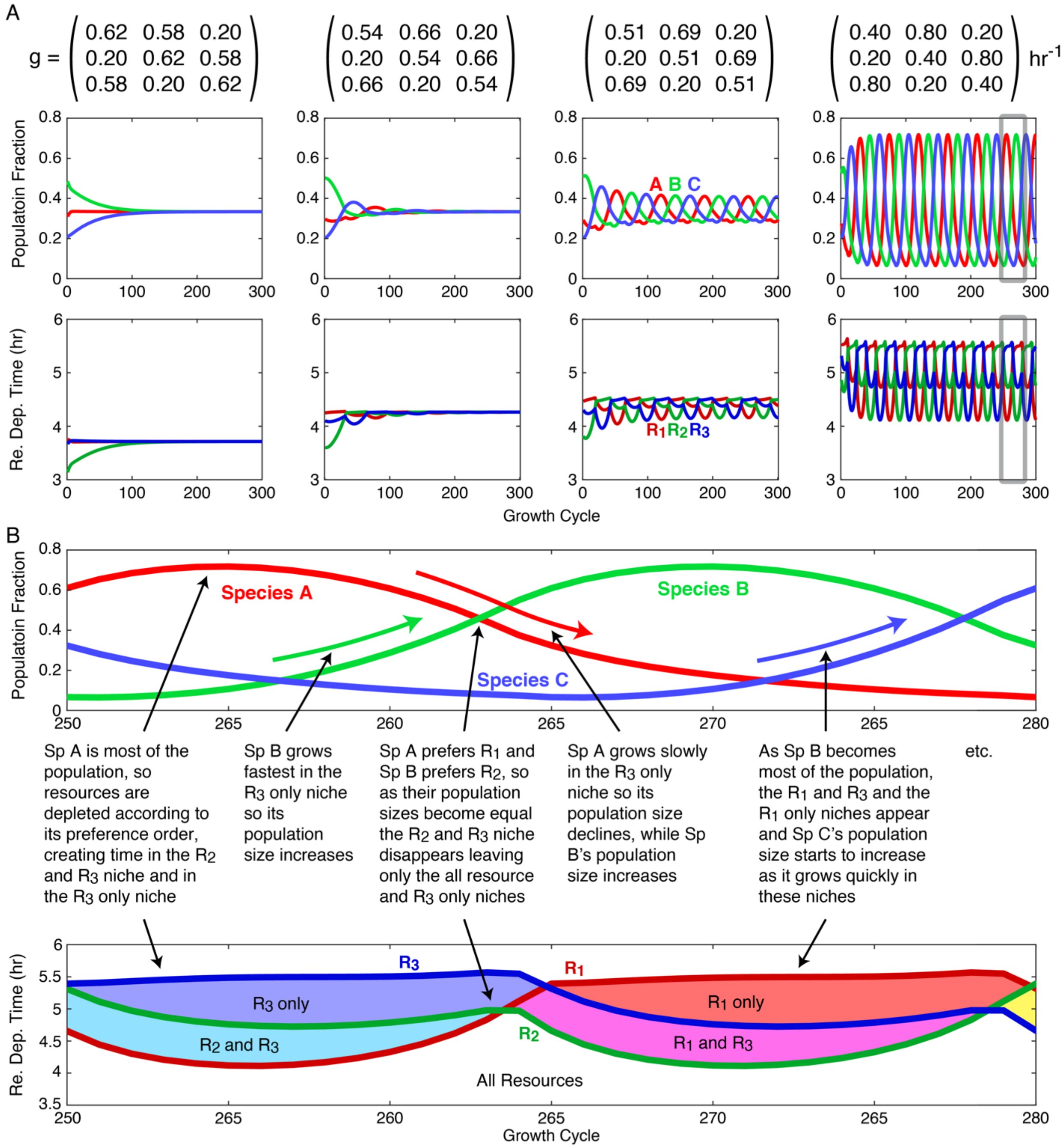


**Appendix Figure S11. Example of a simple constructed oscillation.** (**A**) For species A preferring R1 then R2 then R3, B preferring R2 then R3 then R1, and C preferring R3 then R1 then R2, population fractions at the end of each growth cycle (middle) and resource depletion times on each growth cycle (bottom) as the species' growth rates are varied (top). (**B**) Zoom in on 30 growth cycles from the last (furthest right) growth rate matrix in **A**, annotated with explanations of the interplay between population sizes and resource depletion times.

*§3.6.2 Adding More Species than Resources to the Oscillation*

To add additional species to the oscillation, we can begin by considering the current matrix $\widetilde{\mathbf{G}}$ of growth rates by temporal niche (using the previous species with δg=0.2hr$^{-1}$):

| | **Growth Rates by Temporal Niche (hr$^{-1}$)** | | | | | | |
|---|---|---|---|---|---|---|---|
| | **All Re.** | **$R_1$ & $R_2$** | **$R_1$ & $R_3$** | **$R_2$ & $R_3$** | **$R_1$** | **$R_2$** | **$R_3$** |
| **Sp. A** | 0.4 | 0.4 | 0.4 | **0.8** | 0.4 | **0.8** | 0.2 |
| **Sp. B** | 0.4 | 0.4 | **0.8** | 0.4 | 0.2 | 0.4 | **0.8** |
| **Sp. C** | 0.4 | **0.8** | 0.4 | 0.4 | **0.8** | 0.2 | 0.4 |

Looking at this matrix we observe that each species has two temporal niches in which it is the fastest. If we wanted six species that each had a temporal niche in which they were the fastest growing species (which is not a requirement for coexistence but does make it easier to manually craft examples), we could introduce three species that each grew very quickly on their last preferences. After doing this species A, B, and C will each be the fastest-growing species in one of the two-resource niches and the new species D, E, and F will each be the fastest-growing in one of the single-resource niches. A simulation of these six species community is shown in Appendix Figure S12.

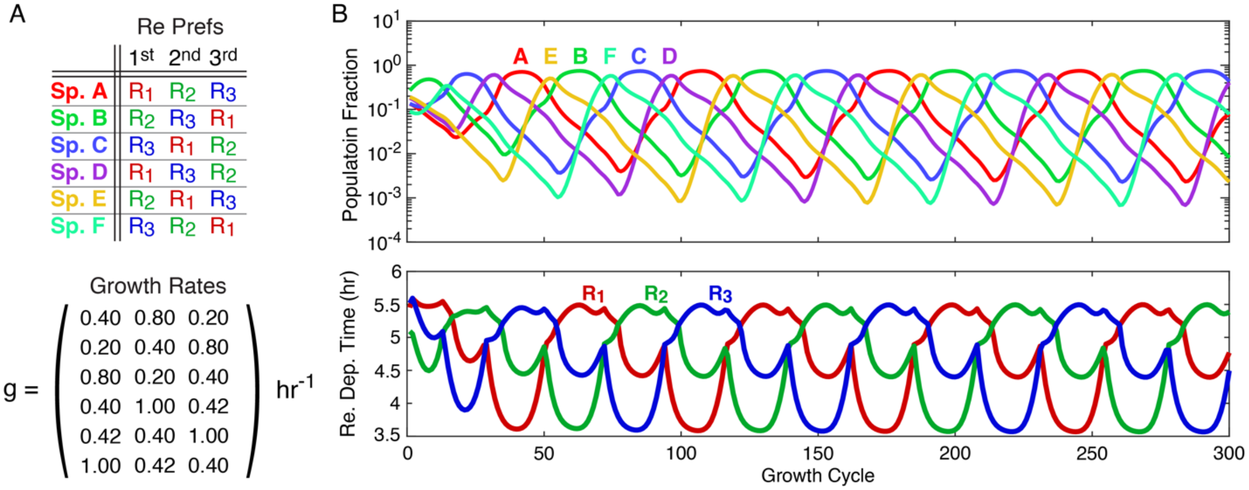


**Appendix Figure S12. Example of a simple constructed oscillation with more coexisting species than resources.** (**A**) Resource preferences and growth rates for the six coexisting species. (**B**) Population fractions at the end of each growth cycle and resource depletion times for these six species competing for the three resources supplied in equal amounts.

*§3.6.3 Constructing an Oscillation on More Than Three Resources*

The approach to constructing a three-species three-resource oscillation with cyclically permuted species (as described in §3.6.1) can be easily extended to more than three resources with species that grow at a medium speed on their first preference, very fast on their second preferences, and slowly on all the rest. With cyclically permuted species (e.g. species A preferring R1 then R2 then R3, species B preferring R2 then R3 then R4, etc.) an oscillation can be easily created, but if the goal is to later add additional species to maximize diversity this is not the best approach as the cyclical structure will prevent all temporal niches from being realized.

An alternative approach is to follow the previous construction more loosely and begin with one species per resource with that species having (i) the resource as its top preference, (ii) all other preferences sampled randomly, (iii) a medium growth rate on its first (and optionally also second) preference, (iv) fast growth rates on its next preference or two, and (v) slow growth rates on all remaining preferences. Randomly sampling communities like this will yield a large variety of oscillations some of which will produce all or nearly all the possible temporal niches. To produce all possible temporal niches, it is important that each resource is the top preference of at least one species or else it can never be the first to be depleted. This form of random sampling is how we produced many of our example communities.

A third approach is two start by combining two communities each competing for a different subset of the supplied resources. Appendix Figure S13 shows two communities based on the discussion in §3.6.1 that oscillate with different periods but similar amplitudes.

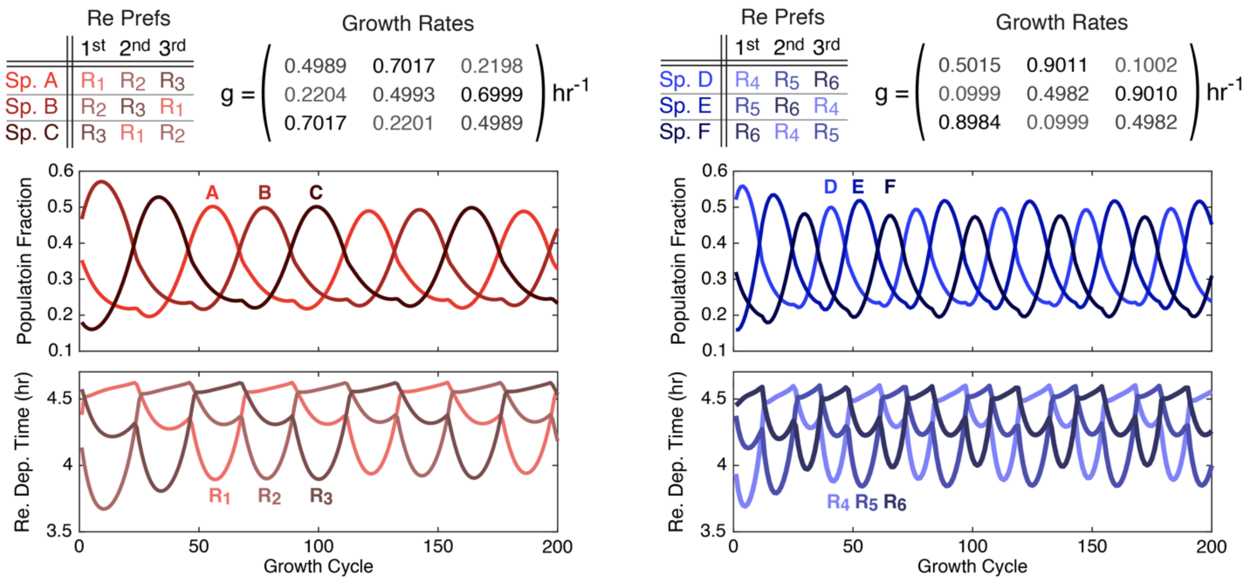


**Appendix Figure S13.** Two example three-species three-resource oscillations based on the discussion in §3.6.1 that oscillate with different periods. A small amount of noise was added to growth rate values after creating species that were exact cyclic permutations of each other to demonstrate that the growth rates do not need to be fine-tuned for these oscillation to occur.

To construct a community competing for six resources, we can begin with the two communities shows in Appendix Figure S13 and have the species A, B, and C primarily compete for R1, R2, and R3 with R4, R5, and R6 as their last preferences and their R4, R5, and R6 growth rates being nearly zero while the species D, E, and F primarily compete for R4, R5, and R6 with R1, R2, and R3 as their last preferences and their R1, R2, and R3 growth rates being nearly zero (Appendix Figure S14A). This creates two essentially independent oscillations (Appendix Figure S14B).

Because the oscillation of A, B, and C competing for R1, R2, and R3 and the oscillation of D, E, and F competing for R4, R5, and R6 had similar amplitudes (when looking at the resource depletion times) but different periods the resource depletion order varies considerably (Appendix Figure S14B) and 58 out of 63 temporal niches are realized (Appendix Figure S14C).

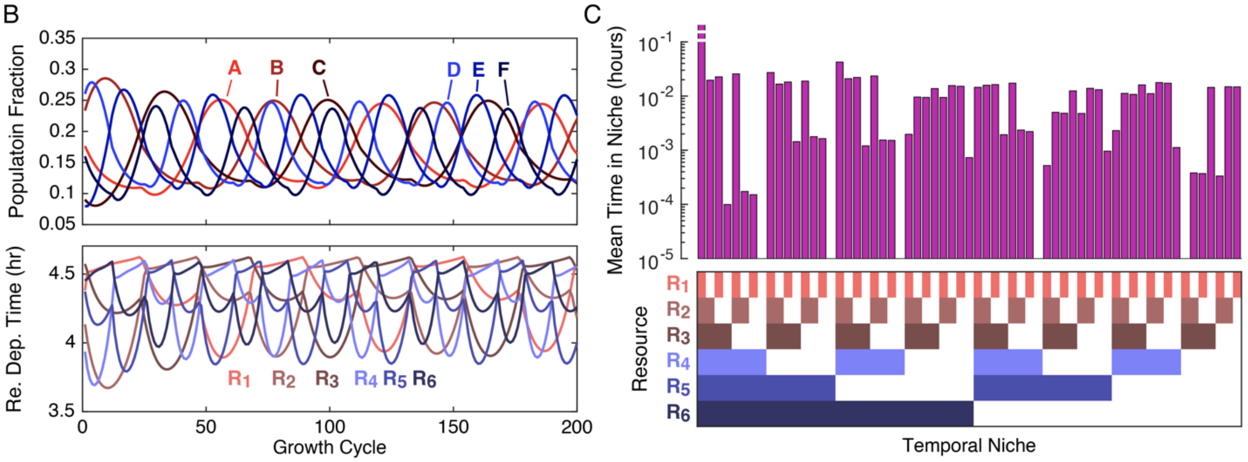


| | 1st | 2nd | 3rd | 4th | 5th | 6th |
|---|---|---|---|---|---|---|
| Sp. A | R1 | R2 | R3 | R4 | R5 | R6 |
| Sp. B | R2 | R3 | R1 | R4 | R5 | R6 |
| Sp. C | R3 | R1 | R2 | R4 | R5 | R6 |
| Sp. D | R4 | R5 | R6 | R1 | R2 | R3 |
| Sp. E | R5 | R6 | R4 | R1 | R2 | R3 |
| Sp. F | R6 | R4 | R5 | R1 | R2 | R3 |

$$g = \begin{pmatrix} 0.4989 & 0.7017 & 0.2198 & 0.0001 & 0.0001 & 0.0001 \\ 0.2204 & 0.4993 & 0.6999 & 0.0001 & 0.0001 & 0.0001 \\ 0.7017 & 0.2201 & 0.4989 & 0.0001 & 0.0001 & 0.0001 \\ 0.0001 & 0.0001 & 0.0001 & 0.5015 & 0.9011 & 0.1002 \\ 0.0001 & 0.0001 & 0.0001 & 0.0999 & 0.4982 & 0.9010 \\ 0.0001 & 0.0001 & 0.0001 & 0.8984 & 0.0999 & 0.4982 \end{pmatrix} hr^{-1}$$

**Appendix Figure S14. Example of a six-species six-resource oscillation constructed from two three-species three-resource oscillations that produces almost all the possible temporal niches.** (**A**) Resource preference orders and growth rates. Note the near independence of A, B, and C competing for R1, R2, and R3 and D, E, and F competing for R4, R5, R6. (**B**) Population fractions and resource depletion times over 200 growth cycles. (**C**) Time spent in each temporal niche. Bar plot on top shows the mean time spent in each temporal niche at steady state. Grid on the bottom defines the temporal niches with a colored box indicating that the resource is present in the niche that the bar directly above corresponds to.

*§3.6.4 General Methodology for Adding Additional Species*

We assume we've found a set of species α through μ that oscillate and we want to add another new species ν to the oscillation. Species ν will consume resources and therefore affect resource depletion times and time spent in each temporal niche at least somewhat, but if we are looking for a species that can join the community without breaking the oscillation or causing any of the original species to go extinct then we can assume an appropriately selected species ν will only change the time spent in each temporal niche by some small amount. We can therefore sample our new species ν by (i) randomly sampling its resource preference order, (ii) sampling its growth rates under the constraint

$$\sum_{\mathrm{tn}} \widetilde{\mathbf{G}}_{\nu,\mathrm{tn}} \langle t_{\mathrm{tn}} \rangle_{\alpha\ldots\mu} = \log(\mathrm{DF}) \,,$$

(where $\widetilde{\mathbf{G}}_{\nu,\mathrm{tn}}$ is the new species' growth rates by temporal niche and $\langle t_{\mathrm{tn}} \rangle_{\alpha\ldots\mu}$ is the mean time spent in each temporal niche before its introduction), and (iii) adding a small amount of random noise to its growth rates (to ensure we're not accidentally creating a fine-tuned, structurally unstable example). Appendix Figure S16 shows an example of adding a new species G to the community presented in Appendix Figure S15 according to this procedure. This example species G happens to be a "medium-at-everything" species with similar growth rates on all resources, but if the goal is to iteratively add as many species as possible it is best to avoid introducing too many of these species (and instead ensure all species have a mix of fast and slow growth rates) or else the oscillations tend to become delicate and sensitive to increasingly small parameter perturbations.

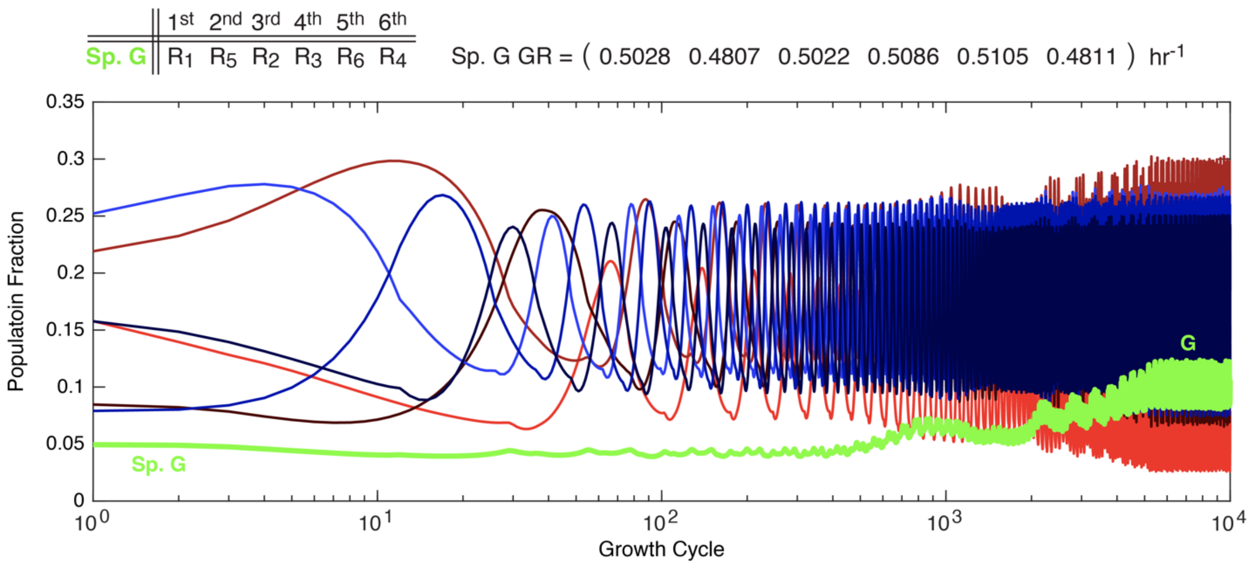


**Appendix Figure S15.** Simulation of the same community as in Appendix Figure S14 but with Species G now added.

#### §3.7 Robustness Against Demographic Noise

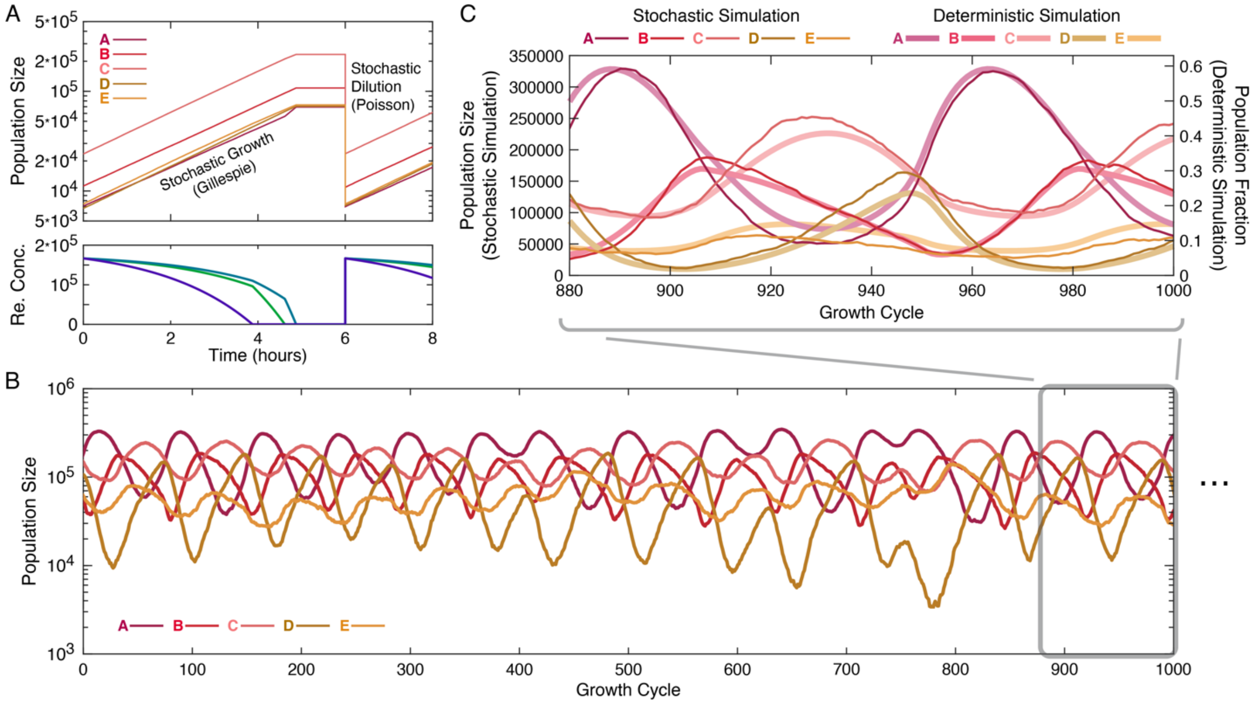


**Appendix Figure S16. Oscillations and enhanced biodiversity are robust against demographic noise.** (**A**) Illustration of our stochastic variation on the Main Text diauxie model. Growth was simulated using a Gillespie algorithm in which (i) a time step was sampled from an exponential distribution with $\langle dt \rangle = 1/\Sigma_\mu g_\mu(t) n_\mu(t)$ where $g_\mu(t)$ is species μ's current per capita growth rate, (ii) a species was randomly sampled proportional to $g_\mu(t) n_\mu(t)$ to have its population size increased by 1, and (iii) resources concentration where updated by $\Delta c_i = -\Sigma_{\mu \text{ eating } \mathrm{R}_i} g_\mu(t) n_\mu(t) dt$. Dilutions were implemented by sampling post-dilution population sizes from Poisson distributions with $\langle n_{\mu,\text{post}} \rangle = \frac{n_{\mu,\text{pre}}}{\text{DF}}$ with $\text{DF} = 10$. Resources were supplied at $c_i(0) = K/3$ with $K = 500000$. (**B**) Population sizes at the end of each of 1000 growth cycles for the community in Main Text Figures 1C and 2 simulated according to the model illustrated and described in **A**. The community sometimes diverges from the limit cycle (for example around growth cycles 700 through 800) but appears to consistently recovery. (**C**) Zoom in of the population sizes at the end of growth cycles 880 through 1000. Thin lines show population sizes in the stochastic simulation. Thicker, paler lines show population fractions in the deterministic simulation for comparison. Despite the moderate deviation from the limit cycle having occurred ~100 growth cycles earlier, the stochastic simulation appears to still closely track the deterministic. Deterministic simulation has a visually tuned time offset to align the two simulations, but both have the same time scale and relative y-axis scales are set according to the carrying capacity $K$ used in the stochastic simulations.

### §4 Additional Results with a Variable Resource Supply

#### §4.1 Competitive exclusion violations are a likely outcome even without metabolic constraints

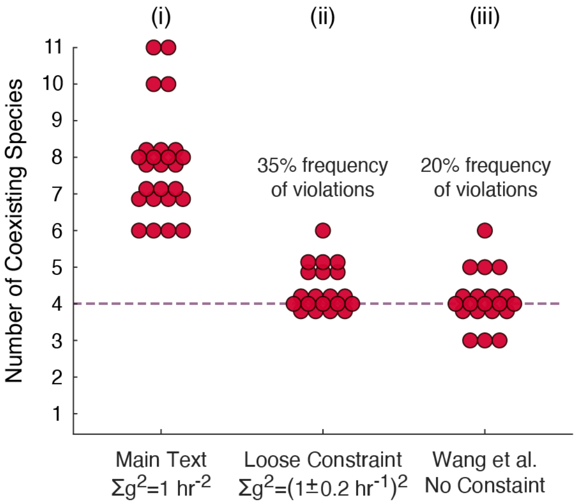


**Appendix Figure S17. Strict metabolic constraints are not necessary for competitive exclusion violations to be a likely outcome.** (**i**) The results on number of surviving species under a fluctuating resource supply from Main Text Figure 4D in which species competing for four resources were sampled from the hypersurface defined by $\sum_{i=1}^{4} {g_{\mu i}}^2 = 1\,\mathrm{hr}^{-2}$, reproduced here to facilitate comparison. (**ii**) To confirm the strict metabolic tradeoffs was not necessary for the competitive exclusion violation, we sampled species using a loose metabolic constraint in which after sampling from the same hypersurface species growth rates were transformed by $g_{\mu i} \to g_{\mu i} * N_\mu(1, 0.2)$ in which $N_\mu(1, 0.2)$ is a normally distributed value sampled once for each species. This meant best-at-everything species were still rare but some species could grow faster on all resources than some of their competitors. Over one third of communities still violated competitive exclusion. (**iii**) We also tested the sampling distribution used in Wang et al. "Complementary resource preferences spontaneously emerge in diauxic microbial communities" Nature Communications 2021. This distribution sampled each component $g_{\mu i}$ independently from $N(0.25\,\mathrm{hr}^{-1}, 0.05\,\mathrm{hr}^{-1})$, thus creating species that tended to all have similar growth rates on all resources with no inverse correlations. Even with no metabolic tradeoffs at all, we still observed a 20% frequency of violations.

#### §4.2 Competitive exclusion violations are a likely outcome even if some resources support universally faster growth rates than others

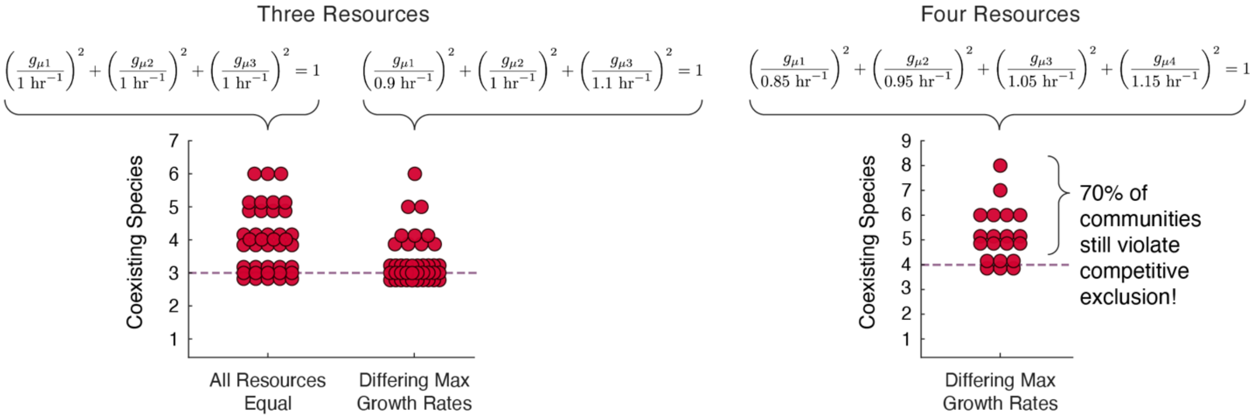


**Appendix Figure S18. Strict equality of resources is not necessary for competitive exclusion violations to be the most likely outcome.** To test whether the exact equal of resources (i.e. that they all supported the same growth rates) we had assumed in the main text was necessary for the diversity we had observed, we repeated the three- and four-resource simulations from Main Text Figure 4D, this time multiplying all $R_1$ growth rates by 0.9, $R_2$ growth rates by 1, and $R_3$ growth rates by 1.1 in the three-resource case and multiplying growth rats by 0.85, 0.95, 1.05, and 1.15 in the four-resource case after sampling from the $\sum_{i=1}^{N_{\mathrm{Re}}} {g_{\mu i}}^2 = 1\ \mathrm{hr}^{-2}$ hypersurfaces. Top row displays the results constraints, showing first the original three-resource constraint (left), then the modify three-resource constraints (middle), and then the modified four-resource constraint (right). In the bottom row, the left plot shows the number of survivors in the Main Text three-resource simulations (left) in comparison to the number of survivors in the three-resources simulations with these modified growth rates (right). The right plot shows the number of survivors in the four-resources simulations with these modified growth rates. Diversity remained high in both the three- and four-resource cases. Even with $R_4$ supporting growth rates 35% faster than $R_1$ supported, 70% of the random communities competing for four resources violated competitive exclusion.

##### §4.3 Sampling from evidence-based resource preference order distributions

A recent paper (Takano, Vila, Miyazaki, Sánchez, and Bajić "The Architecture of Metabolic Networks Constrains the Evolution of Microbial Resource Hierarchies" Molecular Biology and Evolution 2023) explored how the basic structure of central metabolism, specifically which metabolic reactions are and are not possible based on a universal bacterial model, limits which research preference orders are most likely to occur. Takano et al found that certain resource preference orders are vastly more likely to occur than others with some orders being essentially impossible.

The results of our paper depend on species having different resource preference orders, so checking whether our results are robust to enforcing the resource preference orders observed in Takano 2023 is an essential control. In this section, we perform control simulations and observed that, while biodiversity is decreased, communities with more coexisting species than resources are still a frequent occurrence and for certain combinations of resources mean biodiversity is indistinguishable from when resource preference orders are uniformly sampled (Appendix Fig S19).

Takano 2023 includes, amongst other data, the results of random walk simulations in which metabolic reactions were randomly added and removed using a universal database of bacterial metabolism. Takano used flux balance analysis of the resulting metabolic networks to infer likely resource preference orders for a set of seven simple sugars and presents the fifteen most common resource preference orders along with their relative frequencies across the different random walks (Figure 1E from Takano 2023).[1] About one third of random walks resulted in one of these fifteen preference orders (out of 5040 possible preference orders for seven resources) and ordering of resources was highly correlated across these top preference orders (Figure 1E from Takano 2023 and our Appendix Figure S19A). If preference orders between species are indeed as heavily correlated as Takano's results suggest, it is initially unclear whether the increased biodiversity we observed will remain likely as it does depend on at least some variation in preference orders.

To test if we would still see highly diverse communities under the distribution of resource preference orders observed by Takano et al, we performed additional control simulations. We iterated over each subset of three of the seven focal resources in Takano 2023 and calculated the relative frequencies of the three-resource preference orders from the seven-resource preference order frequencies (Appendix Figure S19B). For each subset of resources, we sampled 10 communities of 500 species each with growth rates uniform-randomly sampled from $\Sigma_i g_{\mu i}^2 = 1\,\mathrm{hr}^{-2}$ (as in the Main Text) and ordered to match each species' resource preference order (such that each species' fastest randomly sampled growth rate was assigned to its most preferred resource and its second fastest to its second preference). Communities were simulated in a fluctuating environment with resource

[1] The tenth most frequent preference order is presented with both glucose and mannose as the second preference. For our analysis we removed this preference order and equally distributed its frequency across the preference order with glucose preferred over mannose and the order with mannose preferred over glucose, both of which were already present in the top fifteen preference orders, leaving us with fourteen orders.

supplied uniform-randomly sampled on each growth cycle ($\sigma_{RS} = 0.236$) and the number of coexisting survivors was tallied and explored according to various breakouts (Appendix Fig S19C-F).

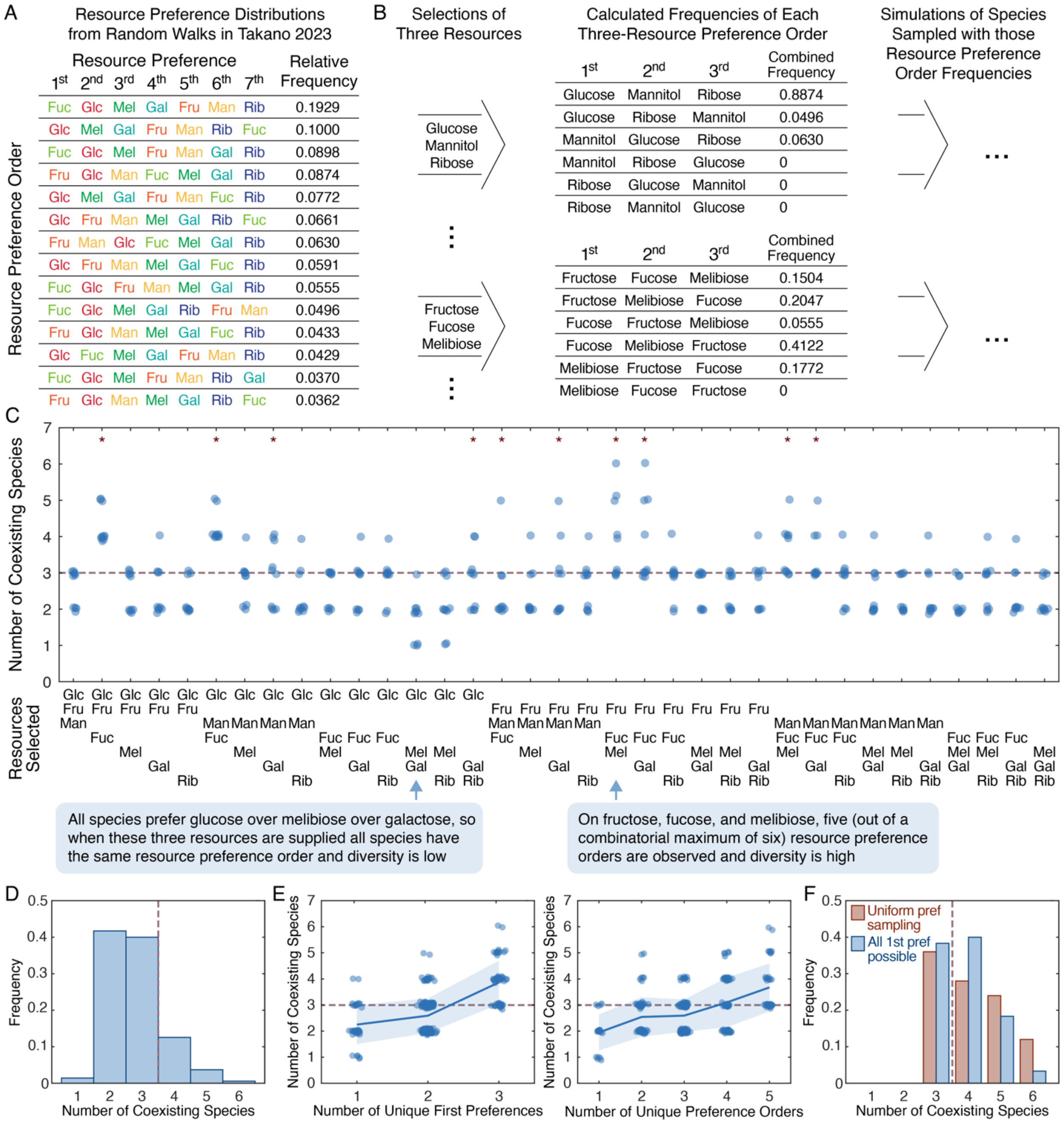


**Appendix Figure S19. Highly diverse communities are possible when species have resource preference orders sampled from an experimentally evidenced distribution, but their likelihood depends on which resources are supplied.** (**A**) Relatively frequencies of the fourteen[1] most frequent resource preference orders as extracted from Figure 1E of Takano 2023 by measuring bar heights. (**B**) Overview of how Takano's results were incorporated into species sampling. We iterated over each subset of three of the seven resources and calculated the frequencies of each of the three-resource preference orders from the frequencies of the seven-resource preference orders. We then sampled communities of 500 species with growth rates sampled from $\Sigma_i g_{\mu i}^2 = 1\ \text{hr}^{-2}$ and then sorted to match each species' resource preference order and simulated them in an environment containing an equal supply of the three resources. (**C**) Number of coexisting species on each subset of three resources. Each point is the result for a separate community of 500 random species. Stars mark three-resource combinations on which mean diversity was not distinguishable at p = 0.05 from mean diversity when resource preferences were uniformly sampled (as in Main Text Figure 4D). Ordered only by which resources are present (bottom). (**D**) Histogram of community diversity across all three-resource combinations. (**E**) Number of coexisting species across all three-resource combinations grouped first by the number of first preferences that were possible given the table in **A** and then grouped by the number of unique resource preference orders. (**F**) Histogram of the number of coexisting species when all three first preferences were possible (blue) and when resource preference were uniformly sampled (red; same data as in Main Text Figure 4D).

We observed that across all ten random communities for each of the 35 three-resource combinations, 17% of random communities featured more coexisting species than resources (Appendix Fig S19D). This result varied considerably based on the identity of the resources. For example, on glucose, melibiose, and galactose only a single of the ten communities even had as many surviving species as resources (Appendix Fig S19C). Meanwhile, on the combination of glucose, fructose, and fucose or the combination of glucose, mannose, and fucose all ten communities featured more coexisting species than resources (Appendix Fig S19C). For 10 of the 35 three-resource combinations, the mean number of surviving species was statistically indistinguishable from the mean number of survivors when resource preference order were uniformly sampled (Appendix Fig S19C).

The variation in how many species each combination of resource typically supports appears, as expected, to be largely driven by how many unique resource preference orders are possible for the combination resources (Appendix Fig S19E). For example, in all the top seven-resource preference orders fructose is preferred over fucose and both those resources are preferred over melibiose, so when these three resources are supplied all species share the same resource preference order (Appendix Fig S19C), which means resources will always be depleted in this order regardless of supply and the full temporal niche structure can never be realized, thus explaining the low diversity of the resulting communities. For other combinations of resources as many as 5 out of 6 possible resource preference orders occurred (Appendix Fig S19B-C) and in these cases diversity was high with 43% of communities having more coexisting survivors than supplied resources (Appendix Fig S19C,E). In order to realize all temporal niches, it is necessary to have at least one species with each resource as its top preference. The number of unique first preferences is therefore a strong predictor of the number of coexisting species, both in theory and in the results (Appendix Fig S19E). Indeed, when all first preferences of possible, observed diversity in the communities based off Takano's results is only barely lower than diversity in the Main Text communities in which resource preferences were uniformly sampled (Appendix Fig S19F).

Thus, highly diverse communities are possible given realistic distributions of resource preference orders but their frequency depends on which specific resource are supplied.

### §5 Seasonal Environmental Cycles

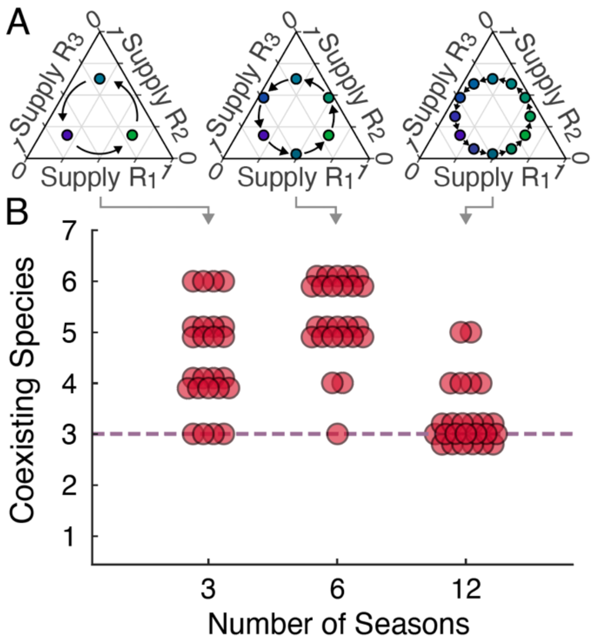


**Appendix Figure S20. Observed community diversity under a periodic environmental oscillations matches expectations.** (**A**) Illustration of three periodic environmental oscillations as defined in Methods using the same resource supply simplex plots as used through the paper. The "three-season" oscillation (left) is expected to produce moderate to high diversity as the large change in resource supply on each growth cycle will prevent the three most specialized species from excluding all those (as would happen under constant environmental conditions) because no ratio of those species could depleted the resources at exactly the same time and the full array of temporal niches is expected to be realized. The "six-season" oscillation (center) is expected to produce even higher diversity due to the addition of growth cycles on which one resource is supplied in a very small amount, which significantly decreases the length of the all-resources-present temporal niche incentivizing generalization into the other niches, thus separating optimal strategies that shared a first preference and increasing the likelihood that all six will be realized in the final community. The "twelve-season" oscillation (right), however, is expected to produce less diverse communities because, while three highly generalized species cannot cause the resources to all be depleted at exactly the same time, the relatively same change in environmental conditions from one cycle to the next allows their population sizes to shift quickly enough to cause resources to be depleted at very similar times, thus disincentivizing investment in resources other than one's top preference. (**B**) Observed diversity of the final communities after sampling 500 species uniform randomly from $\sum_i g_{\mu i}^2 = 1\ \mathrm{hr}^{-2}$, setting resource preferences equal to species' descending growth rate orderings, running the simulation for $1.2*10^6$ growth cycles, and visually checking for and manually removing any species that were clearly going slowly extinct. The observed diversity closely matches the predictions.

### §6 Resource Preference Complementarity and Anomalous Species

A recent report (Wang, Goyal, Dubinkina, George, Wang, Fridman, and Maslow "Complementary resource preferences spontaneously emerge in diauxic microbial communities" *Nature Communications* 2021) demonstrated that within our model of diauxic resource competition under a constant resource supply (i) survivors almost always had "complementary" resource preferences (such that there was usually exactly one survivor with each resource as its top preference) and (ii) that anomalous species (whose resource preference order did not match their ordering of their growth rates) rarely survived. Here we test whether our final communities of coexisting species also share these properties and confirm that they do.

Because our final communities typically featured more surviving species than resources, multiple species shared first (and even first few) resource preferences, thus prompting a definition of complementary that would be equivalent with Wang 2021 when considering first preferences and only as many species as resources but also extend nicely to higher preferences and larger communities. To this end, resource preference complementarity was calculated for the communities whose number of survivors are shown in Figure 4D as:

$$\text{Complimentary of N}^{\text{th}}\text{ preferences amongst species sharing 1}^{\text{th}}\text{ through }(\text{N}-1)^{\text{th}}\text{ preferences}$$
$$= \frac{(\text{Number of unique N}^{\text{th}}\text{ preferences those species have})}{\min[(\text{Number of species in that set}), (\text{Number of possible N}^{\text{th}}\text{ preferences})]}$$

This calculation generally produced multiple data points for each community at each order of preference as it considered the groups of species with the same earlier preferences separately.

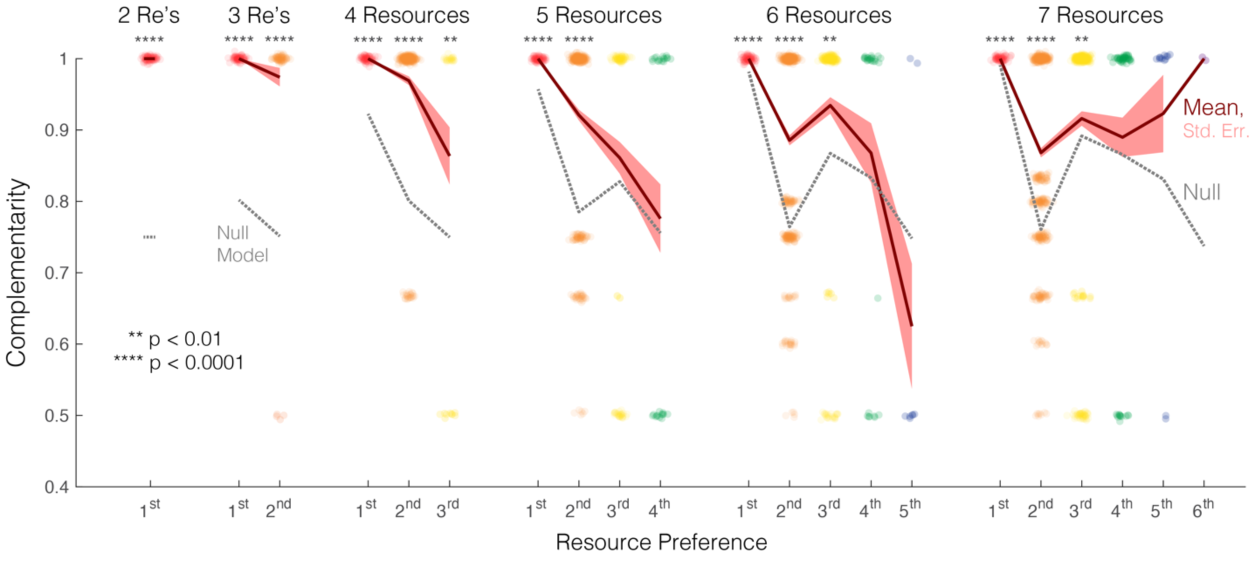


**Appendix Figure S21. Complementary of resource preferences.** Complementarity of resource preferences using the above-defined formula by number of resources in the simulation and resource preference ordinal. Null model is calculated from taking the number of surviving in each community and calculating the expected complementarities if all species had randomly assigned resource preferences.

Complementarity was observed to indeed be high. For example, every final community had each first preference represented by at least one species – representing full first order complementarity. Similarly, 65.3% of the sets of coexisting species sharing first preferences and 81.0% of the sets of coexisting species sharing first and second preferences were fully complementary (either all having different next preferences or having all remaining resources represented as next preferences).

We additionally tested whether our final communities, like those produced in Wang 2021, had anomalous species rarely surviving. This test was done both for comparison to the previous report and to rule out the presence of anomalous species as essential to our results. Our Main Text simulations were run in two batches, one with anomalous species included and one without. Specifically, when we were varying the magnitude of environmental fluctuations in three-resource environments (Main Text Figures 4C and 5) we included anomalous species, and when we were varying the number of resources (Main Text Figure 4D) we excluded anomalous species. We show in Appendix Figure S22 that anomalous species very rarely survived (and only did when the out-of-order growth rates were very close anyways). Their inclusion, therefore, should not have had any significant impact on our results.

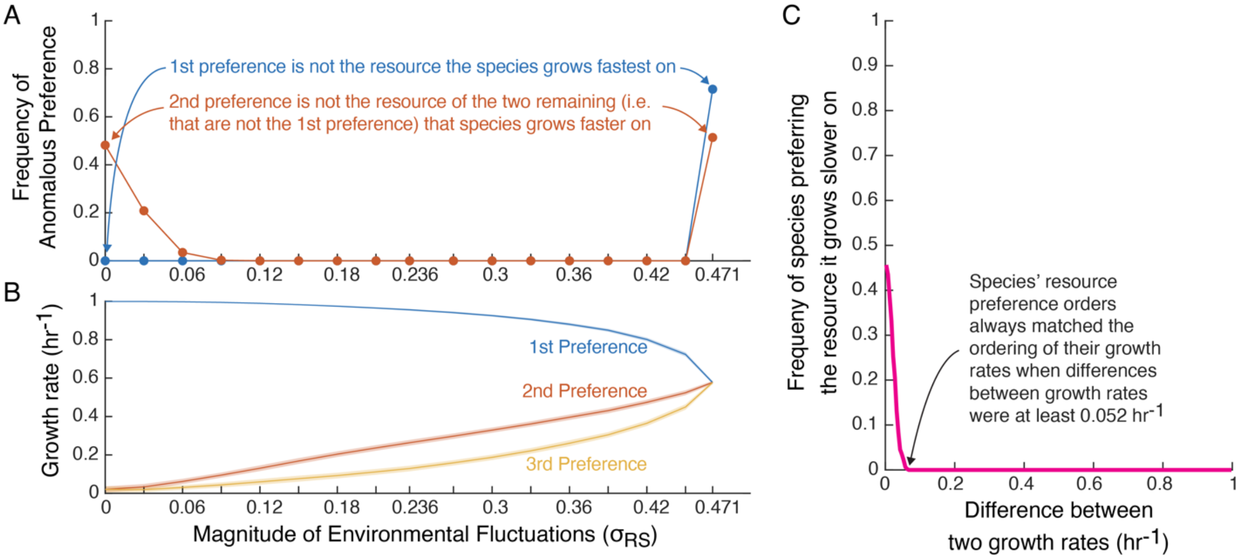


**Appendix Figure S22. Anomalous species rarely survived.** (**A**) Frequency at which a surviving species did not have the resource it grew fastest on as its top preference (blue) and at which it grew faster on its third preference than its second preference. By fluctuation magnitude and using results from the same simulations as in Main Text Figures 4C and 5. (**B**) For comparison the mean growth rates of surviving species by resource preference. Shaded regions indicate one standard deviation. (**C**) Frequency of a species preferring one resource over another that it grows slowest on as a function of the difference between the two growth rates.

#### §7 Comparison to a Model Without Sequential Resource Utilization

To confirm and further explore the importance of sequential resource utilization specifically to enabling highly diverse communities, we here explore a simple variation on the Main Text diauxie model that removes sequential utilization while leaving all other dynamics as intact as possible. We demonstrate that with sequential utilization removed highly diverse communities become impossible.

The growth dynamics in the Main Text model can be thought of as having two main features: (i) discrete resource utilization states with constant growth rates such that $\frac{\partial}{\partial c_i}\left(\frac{dn_\mu}{dt}\right) = 0$ for $c_i > 0$ and (ii) increases in the consumption in one resource after another is depleted. The latter feature is the notion of sequential utilization.

Perhaps the easiest way to remove the feature of sequential resource utilization without changing any other model dynamics would be to have species each consume a single resource until it is depleted and then stop growing. This modification would create relatively uninteresting dynamics in which the species competing for resource $\mathrm{R}_i$ and those competing resource $\mathrm{R}_{j\neq i}$ do not interact and each resource independently supports whichever species has the fastest growth rate on it with no mechanisms for additional coexistence. We therefore look for a slightly more interesting model to explore.

To remove sequential utilization while still having interesting, nontrivial dynamics that closely resemble those of the Main Text diauxie model, we can define species as initially coutilizing resources (with constant growth and per capita resource uptake rates) and consuming an increasingly small subset of resources as depletions occur. Importantly, species do not increase their uptake of resources or how much growth they gain from them as other resources run out. Mathematically, this model would be defined by

$$n_\mu'(t) = n_\mu(t)\Sigma_i g_{\mu i}\Theta[c_i(t)]$$

and

$$c_\mu'(t) = -\Theta[c_i(t)]\Sigma_\mu g_{\mu i} n_\mu(t)\ ,$$

where $\Theta[c] = 1$ for $c > 0$ and $\Theta[c] = 0$ for $c = 0$ and a periodic dilution and resource resupply occur as in the Main Text model.

As we did in our analysis of the Main Text model, we can ask by how much species' log-populations grow on each growth cycle. This is accomplished by integrating their growth rates over time. Doing so and plugging in the above expression for $n_\mu'(t)$, we obtain

$$\Delta\log[n_\mu] = \int_0^{t_{\text{dep,last}}} \frac{n_\mu'(t)}{n_\mu(t)}dt = \int_0^{t_{\text{dep,last}}} \Sigma_i g_{\mu i}\Theta[c_i(t)]\,dt = \Sigma_i g_{\mu i}\int_0^{t_{\text{dep,last}}} \Theta[c_i(t)]\,dt\ .$$

The integrals of the step functions are just the length of time until the resource concentrations reach zero – i.e. the depletion times – and do not depend on the resource depletion order. Therefore,

$$\Delta\log[n_\mu] = \Sigma_i g_{\mu i} t_{\text{dep},i}\ .$$

The average of a species' log-population growth across many growth cycles is a simple linear function of the average depletion time of each resource, specifically

$$\langle \Delta \log[n_\mu] \rangle = \Sigma_i g_{\mu i} \langle t_{\mathrm{dep},i} \rangle \,.$$

Importantly, nothing in the above calculation depends on the order of $\{t_{\mathrm{dep},i}\}$. This independence arises because nothing changes about the species' consumption of resource $\mathrm{R}_i$ when resource $\mathrm{R}_{j \neq i}$ is depleted.

If we impose that $\langle \Delta \log[n_\mu] \rangle = \mathrm{DF}$ for all surviving species (which is the condition that all species are exactly keeping up with the dilution factor) then by the above equation only have one degree of freedom per resource, it is impossible to have average depletion times $\langle t_{\mathrm{dep},i} \rangle$ that allow more species than resources to coexist. Thus, when we remove sequential resource depletion while leaving all other model dynamics as intact as possible, highly diverse communities become impossible. This contrast to the Main Text diauxie model illustrates the central importance of the specific aspect of sequential resource utilization to the possibility of highly diverse communities. In this modified model, the time spent in each temporal niche does not directly matter. Instead all that impact a species' growth is the total time each resource is present, reducing the relevant set of niches to simply being one niche per resource.

The illustrate this analytic result, we sampled 300 pools of 500 species competing for 3 resources from the same $\Sigma_i g_{\mu i}^2 = 1\ \mathrm{hr}^{-2}$ distribution as in the Main Text and simulated them according to the above-described model in an environment in which the resource supply was uniform-randomly sampled ($\sigma_{\mathrm{RS}} = 0.236$) on each growth cycle and tallied the number of coexisting survivors in each community (Appendix Fig S23B). We compared these tallies to the number of coexisting species in the Main Text diauxie model (Appendix Fig S23A, same data as the $N_{\mathrm{Sp}} = 3$ condition in Main Text Figure 4D) and to the classic linear resource consumption model defined by

$$n'_\mu(t) = n_\mu(t) \Sigma_i g_{\mu i} c_i(t)$$

and

$$c'_\mu(t) = -c_i(t) \Sigma_\mu g_{\mu i} n_\mu(t) \,,$$

which was also simulated under the same periodic dilution and uniform-randomly sampled resource supply (Appendix Fig S23C).

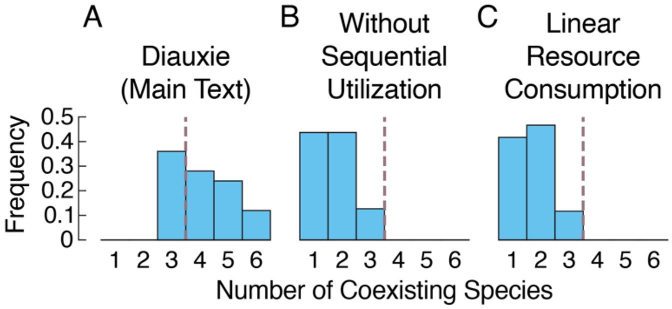


**Appendix Figure S23. When sequential resource utilization is removed, highly diverse communities become impossible.** (**A**) Histogram of the number of coexisting species in the simulations of 500 species competing for three resources supplied in uniform-randomly sampled fractions on each day originally presented in Main Text Figure 4D. (**B**) Histogram of the number of coexisting species in the modified model with sequential resource utilization removed (described in Appendix §7) simulated under the same environmental conditions. (**C**) Histogram of the number of coexisting species in the linear resource consumption model simulated under the same environmental conditions.

As expected, we saw no cases of more coexisting species than supplied resources under either of the models which lacked sequential resource utilization (Appendix Fig S23B-C), in notable contrast to the high biodiversity supported by the Main Text diauxie model (Appendix Fig S23A).

There are, of course, other models which lack sequential resource utilization but nevertheless support more coexisting species than resources (Armstrong 1976, Levins 1979, Huisman 1999, Posfai 2017, Fridman 2022, etc.). Sequential utilization and the temporal niches it creates are one interesting mechanism by which highly diverse communities can emerge, but it is not the only.

This Appendix section demonstrates how removing sequential resource utilization from the Main Text diauxie model and replacing it with simultaneous utilization prevents the stable coexistence of more species than resources. This is because without sequential utilization there is no fundamental reshuffling of which species are in direct competition and for which resources at each resource depletion event. This reshuffling is a fundamental and specific consequence of sequential utilization and the origin of the temporal niche structure that expands the potential for highly diverse communities to coexist. This Appendix exploration thus illustrates the specific importance of sequential utilization to the emergence of highly diverse communities in our Main Text results.
